# The Great Deceiver: miR-2392’s Hidden Role in Driving SARS-CoV-2 Infection

**DOI:** 10.1101/2021.04.23.441024

**Authors:** J. Tyson McDonald, Francisco Javier Enguita, Deanne Taylor, Robert J. Griffin, Waldemar Priebe, Mark R. Emmett, Mohammad M. Sajadi, Anthony D. Harris, Jean Clement, Joseph M. Dybas, Nukhet Aykin-Burns, Joseph W. Guarnieri, Larry N. Singh, Peter Grabham, Stephen B. Baylin, Aliza Yousey, Andrea N. Pearson, Peter M. Corry, Amanda Saravia-Butler, Thomas R. Aunins, Sadhana Sharma, Prashant Nagpal, Cem Meydan, Jonathan Foox, Christopher Mozsary, Bianca Cerqueira, Viktorija Zaksas, Urminder Singh, Eve Syrkin Wurtele, Sylvain V. Costes, Gustavo Gastão Davanzo, Diego Galeano, Alberto Paccanaro, Suzanne L. Meinig, Robert S. Hagan, Natalie M Bowman, UNC COVID-19 Pathobiology Consortium, Matthew C. Wolfgang, Selin Altinok, Nicolae Sapoval, Todd J. Treangen, Pedro M. Moraes-Vieira, Charles Vanderburg, Douglas C. Wallace, Jonathan Schisler, Christopher E. Mason, Anushree Chatterjee, Robert Meller, Afshin Beheshti

## Abstract

MicroRNAs (miRNAs) are small non-coding RNAs involved in post-transcriptional gene regulation that have a major impact on many diseases and provides an exciting avenue towards antiviral therapeutics. From patient transcriptomic data, we have discovered a circulating miRNA, miR-2392, that is directly involved with SARS-CoV-2 machinery during host infection. Specifically, we show that miR-2392 is key in driving downstream suppression of mitochondrial gene expression, increasing inflammation, glycolysis, and hypoxia as well as promoting many symptoms associated with COVID-19 infection. We demonstrate miR-2392 is present in the blood and urine of COVID-19 positive patients, but not detected in COVID-19 negative patients. These findings indicate the potential for developing a novel, minimally invasive, COVID-19 detection method. Lastly, using *in vitro* human and *in vivo* hamster models, we have developed a novel miRNA-based antiviral therapeutic that targets miR-2392, significantly reduces SARS-CoV-2 viability in hamsters and may potentially inhibit a COVID-19 disease state in humans.

## Introduction

In Fall of 2019, the zoonotic spillover event led to the first known human infection with the severe acute respiratory syndrome coronavirus 2 (SARS-CoV-2) and subsequent human-to-human transmission triggered a pandemic leading to a worldwide health crisis from the resulting disease, referred to as coronavirus disease 2019 (COVID-19) (Huang et al., 2020; Zhu et al., 2020). COVID-19 causes substantial pulmonary disease but can also cause systemic health risks from extrapulmonary manifestations. Its effects entangle the entire body including but not limited to the cardiovascular, gastrointestinal, and hematological systems that may lead to long lasting effects after the virus has left the body, known as PASC (post-acute sequela of COVID-19) (Carfi et al., 2020; Feng et al., 2020; Jacobs et al., 2020). SARS-CoV-2 is classified as a member of the Coronaviridae family, viruses with a enveloped positive-stranded RNA with the ability to infect cross-species (V’Kovski et al., 2021). Currently, three vaccines have been approved for emergency use by the Food and Drug Administration (Baden et al., 2021; Polack et al., 2020; Sadoff et al., 2021). While these vaccines represent a favorable milestone, additional data is required to demonstrate their long-term effectiveness against SARS-CoV-2 and protection against new strains. To prevent an endemic, the complete global eradication of COVID-19 will require a wide majority of the world’s population to be vaccinated to achieve herd immunity. Unfortunately, a portion of the population will not get vaccinated. Therefore, strategies for therapeutic options against COVID-19 are particularly relevant and important to explore to treat severe illnesses and overcome this global pandemic. Currently the majority of antivirals are repurposed drugs utilized for other disease and have shown limited clinical efficacy, such as remdesivir (Abdelrahman et al., 2021). This brings a needed urgency to develop antivirals specifically designed against SARS-CoV-2.

A potential avenue for an alternative antiviral agent is to target specific microRNAs (miRNAs) associated with SARS-CoV-2 infection and subsequent manifestation of COVID-19. MicroRNAs (miRNAs) are non-coding RNAs that are involved with regulation of post-transcriptional gene expression and can impact entire pathways related to viruses and diseases (Jiang et al., 2009; Trobaugh and Klimstra, 2017). Each miRNA can target multiple messenger RNAs (mRNAs) and taken together, miRNAs are predicted to regulate over half of the human transcriptome (Friedman et al., 2009). Recent evidence has shown different diseases, including COVID-19, leads to distinct complements of miRNAs in the blood (Mishra et al., 2020; Nersisyan et al., 2020; Sacar Demirci and Adan, 2020; Teodori et al., 2020). These circulating miRNAs are highly stable and have the potential to be used for minimally invasive novel detection, potential biomarkers, and therapeutic targets (Tribolet et al., 2020). The interactions between miRNAs and viruses have revealed a multifaceted relationship. Specifically, viruses have been shown to avoid the immune response by leveraging cellular miRNAs to complete their replication cycle (Trobaugh and Klimstra, 2017). The following mechanisms are central to the interaction of viruses and miRNAs: 1) miRNA processing can be blocked by viruses interacting with key proteins such as Dicer and associated proteins, 2) viruses can sequester miRNAs resulting in dysregulation of specific target mRNAs, 3) viruses can utilize miRNAs to redirect regulatory pathways of other miRNA targets to provide survival advantages, and 4) viruses can directly encode miRNA precursors that are processed by the canonical miRNA cellular pathway and have well-defined functions to specifically target and regulate the viral replicative cycle (Schult et al., 2018; Trobaugh and Klimstra, 2017).

Here, we report on a miRNA, miR-2392, that may directly regulate and drive a COVID-19 response. This miRNA is predicted from COVID-19 RNA-sequencing patient data that resulted in multiple miRNAs being suppressed/inhibited (miR-10, miR-10a-5p, miR-1-3p, miR-34a-5, miR-30c-5p, miR-29b-3p, miR-155-5p, and miR-124-3p) and one miRNA being upregulated (miR-2392). With further examination, we hypothesize miR-2392 to be a key miRNA involved with COVID-19 progression. Specifically, miR-2392 drives downstream suppression of mitochondria activity while increasing inflammation, glycolysis, and hypoxia. MiR-2392 upregulation is concomitant with symptoms associated with COVID-19 infection in the host. Patient blood and urine data show that miR-2392 circulates in COVID-19 infected patients and its concentration increases as a function of viral load. Our results demonstrate that miR-2392 may be utilized as an effective biomarker of COVID-19. Furthermore, we have started development of a miR-2392 inhibitor and provide evidence that its use reduces SARS-COV-2 viability in targeted viral screens with A549 cells and reduces the impact of infection in COVID-19 animal models. With further development this miR-2392 inhibitor may represent an effective antiviral therapeutic towards inhibiting the virus and limiting a negative host response from COVID-19 in humans.

## Results

### Identification of key miRNAs associated with COVID-19 infection

Currently, the majority of publications associated with miRNAs and SARS-CoV-2 are based on *in silico* predictions. To identify miRNAs that may be involved in driving COVID-19 severity in the host, we examined publicly available Bronchial Alveolar Lavage Fluid (BALF) RNA-sequencing (RNA-seq) data from 13 individuals. Differential gene expression was assessed using a 1.2-fold changes for p-values less than 0.01 revealing 42 increased and 347 decreased genes compared to controls. Using the upstream regulator analysis from the Ingenuity Pathway Analysis (IPA) knowledge database, the miRNAs from differentially expressed genes (FDR<0.05) from COVID-19-positive patients were inferred. Eight miRNAs were predicted to drive significant changes in COVID-19 positive patients with the downregulation of seven miRNAs (miR-10, miR-1, miR-34a-5p, miR-30c-5p, miR-29b-3p, miR-124-3p, and miR-155-5p) and upregulation of a single miRNA, miR-2392 (**Fig. 1A**). Using IPA’s downstream effects analysis to predict biological processes from the combined suppression of the seven miRNAs and the upregulation of miR-2392 resulted in increased inflammation, immune suppression, and suppression of mitochondrial activity in the BALF dataset (**Fig. 1B and 1C**).

**Figure 1.**
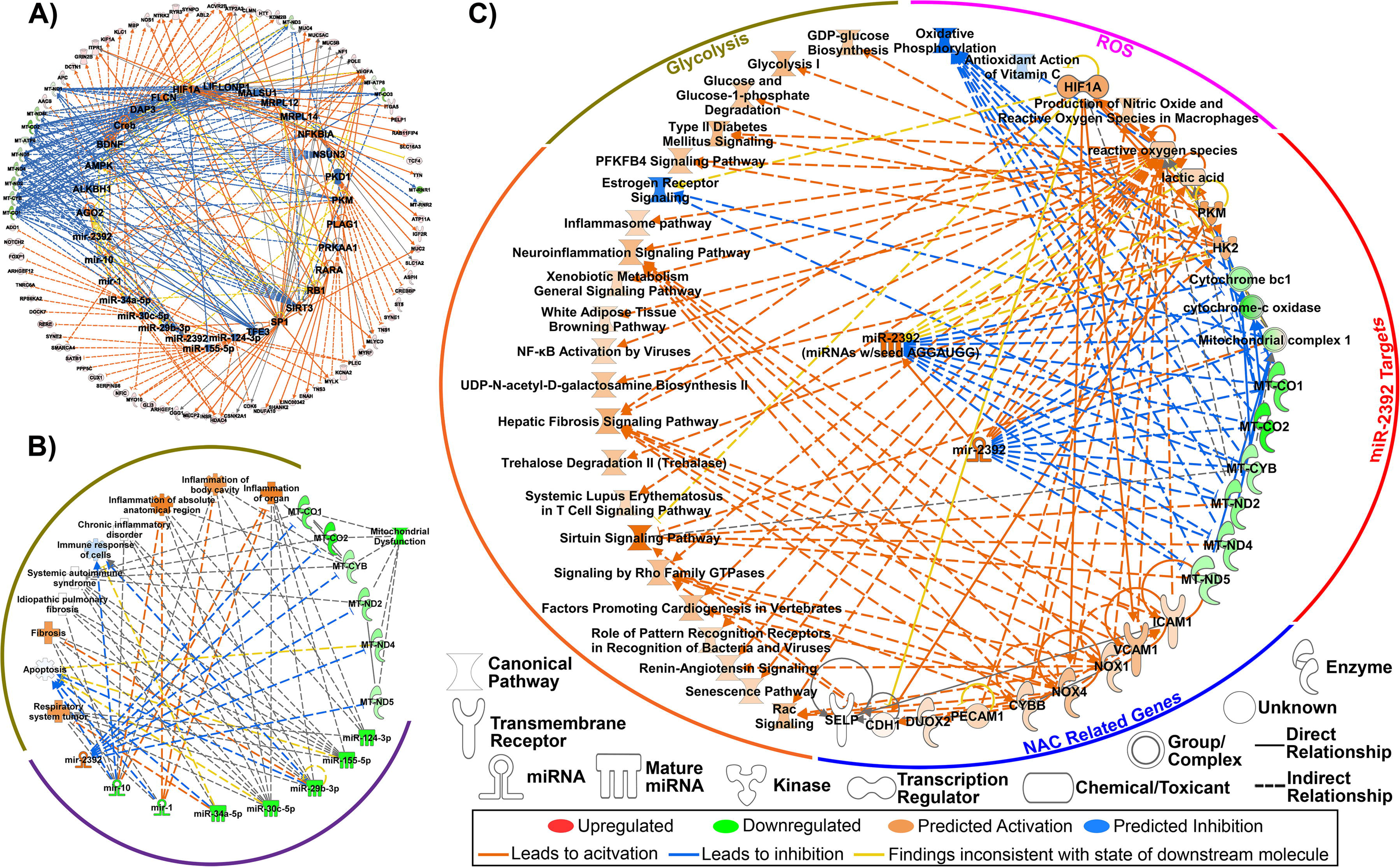
Key miRNA signature as predicted from Bronchial Alveolar Lavage Fluid (BALF) RNA-seq data in patients with COVID-19. **A)** Predicted upstream regulators determined through Ingenuity Pathway Analysis (IPA) consistent with the transcriptional response from differentially expressed genes (FDR<0.05; outer ring). Eight miRNAs were among the key regulators in response to COVID-19 (inner ring). **B)** Major biological responses resulting from dysregulation of this eight miRNA signature drive immune- and inflammatory-related pathways as well as mitochondrial dysfunction determined through IPA. **C)** Pathway regulation by miR-2392 from BALF RNA-seq data determined through IPA.

In support of these findings, previous studies have shown upregulation of miR-10, miR-124, or miR-1 have antiviral roles during infection (Hu et al., 2020; Sardar et al., 2020; Yang et al., 2016) and upregulation of miR-30 and miR-155 causes suppression of other coronaviruses (Dickey et al., 2016; Ma et al., 2018). The one miRNA predicted to be upregulated in COVID-19 patients from the BALF data was miR-2392. Though limited, the existing literature on miR-2392 demonstrates it is related to mitochondrial suppression and increased glycolysis (Fan et al., 2019) and circulating factors related to negative health risks (Chen et al., 2013; Fan et al., 2019; Li et al., 2017; Yang et al., 2019).

We performed pathway analysis with miR-2392 targets to determine its potential impact on the host. Upregulation of miR-2392 in the BALF RNA-seq dataset impacted many downstream targets and pathways related to negative health outcomes (**Fig. 1C**). This includes mitochondrial suppression and activation of factors related to reactive oxygen species (ROS). Since it is known that miR-2392 directly interacts with mitochondrial DNA (mtDNA) to inhibit the levels of oxidative phosphorylation transcripts (Fan et al., 2019), this could be a compensatory response to the inhibition of mitochondrial bioenergetics.

Glycolytic pathways (**Fig. 1C**) are also upregulated in association with increased miR-2392. MiR-2392 drives both hexokinase 2 (HK2) and pyruvate kinase (PKM), both of which positively regulate glycolysis. HK2 produces a primary regulator of glycolysis, glucose-6-phosphate and enhances GDP-glucose biosynthesis. GDP-glucose is a nucleotide sugar which an essential substrate for all glycosylation reactions (i.e. glycosylation of viral spike proteins). PKM is essential for the production of ATP in glycolysis that catalyzes the transfer of the phosphate group from phosphoenolpyruvate to ADP to make ATP. The mechanism of how miR-2392 is driving these pathways is not clearly understood, but one possibility could be stabilization of glycolytic transcripts.

Overall, the observed upregulation of glycolysis and antiviral effects related to miR-2392 suppression are consistent with the recently documented role of glucose metabolism in the progression of viral infection and poor outcome of COVID-19 (Ardestani and Azizi, 2021). It is also consistent with the reported effects of suppression of glycolysis by inhibitors like the glucose analog, 2-deoxy-D-glucose (2-DG), that was shown to suppress *in vitro* SARS-CoV-2 replication (Ardestani and Azizi, 2021; Bojkova et al., 2020). Interestingly, 2-DG is also 2-deoxy-D-mannose and as such can interfere with processes utilizing mannose, a monosaccharide that is produced *in vivo* from glucose. Replacement of a mannose molecule by 2-DG in the respective SARS-CoV-2 N- or O-glycans might lead to their truncation and suppression of virus infectivity and proliferation. These and miR-2392 data indicate targeting of glucose metabolism might have significant impact on SARS-CoV-2 infections.

Targets implicated in antioxidant N-acetyl cysteine (NAC) therapy are to be upregulated (**Fig. 1C**). These include adhesion molecules of activated endothelial cells such as intercellular adhesion molecule 1 (ICAM1), vascular cell adhesion molecule 1 (VCAM1), and E-selectin, which allow attachment of hematopoietic immune and non-immune cells to the endothelial surface, and thus, contribute to inflammation and activation of the coagulation cascade. Powerful antioxidants such as NAC potentially counteract COVID-19 infections by suppressing viral replication via improving intracellular thiol redox ratio as a precursor for major thiol antioxidant glutathione (Ho and Douglas, 1992) and inhibiting the NF-kB pathway (Poppe et al., 2017). Inhibition of the NF-κB pathway reduces inflammatory damage by altering the glutathione and glutathione disulfide ratio (Aykin-Burns et al., 2005; Griffin et al., 2003; Jia et al., 2010). Because NAC can also modulate oxidative burst and reduce cytokine storm without weakening the phagocytizing function of neutrophils (Aykin-Burns et al., 2005; Jia et al., 2010), clinical trials in COVID-19 patients are being conducted (Alamdari et al., 2020) to reduce severe infection (Ibrahim et al., 2020). These results were in line with the role of miR-2392 in reducing the activities of electron transport chain complexes and enhancing glycolysis. It is also speculated that NAC could inhibit binding to ACE2 by reducing disulfide bonds in the receptor-binding domain. Inflammatory pathways and others that are observed with COVID-19 infection were also seen to be activated downstream of miR-2392.

### Conservation of miR-2392 between species and its predicted interactions with the SARS-CoV-2 genome

Viral miRNAs can facilitate interspecies viral transmission due to the high conservation of miRNAs among species and the ability of viruses to integrate miRNAs into its own genome (Sacar Demirci and Adan, 2020; Schult et al., 2018). This has been shown to assist viral replication and evasion of the immune system (Islam and Islam, 2021). Thus, we analyzed the conservation of human miR-2392 across species and the integration of miR-2392 into the SARS-CoV-2 genome (**Fig. 2**).

**Figure 2.**
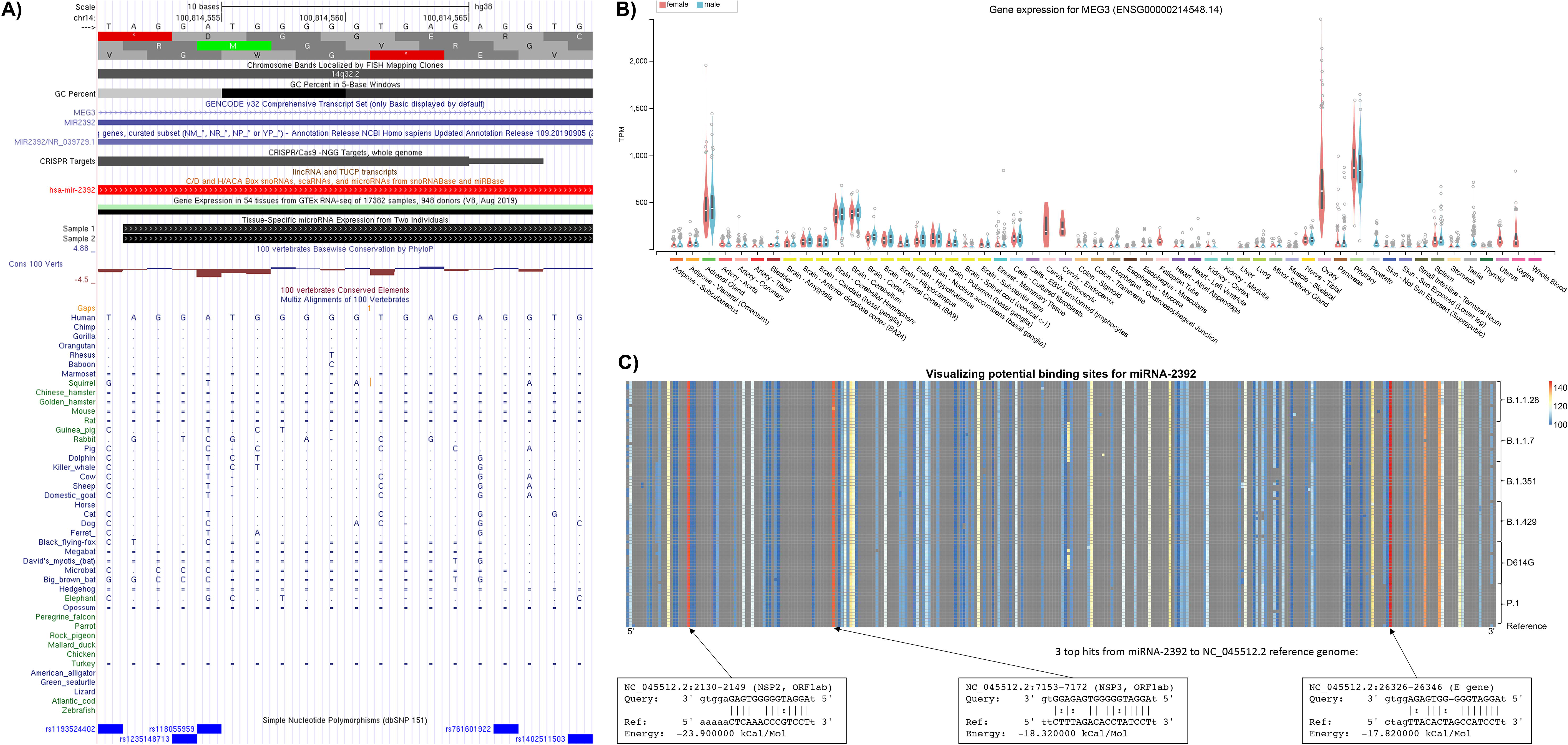
Cross-species and viral integration of miR-2392. **A)** The conservation of miR-2392 across species determined by UCSC Genome Browser. Boxes (▪) represent aligning and conserved sequence regions. Double horizontal line (=) represents both the genome and query have unalignable sequence between regions of aligned sequence, a double-sided insertion. Single lines (-) indicate gaps. **B)** The expression of MEG3, the miR-2392 host gene, in different tissues from healthy patients. **C)** Potential binding sites of miR-2392 visualized across the SARS-CoV-2 genomes (NC045512.2 reference and lineages of concern from GISAID). Average miRanda scores are for hits within the 100bp window. Three top hits are shown.

We determined the conservation of miR-2392 across different species using the UCSC Genome Browser (Kent et al., 2002). The mature 20 base-pair miR-2392 is derived from an 84 base-pair region of the 3’-UTR in the long non-coding RNA gene, maternally expressed 3 (MEG3) and is located in an imprinted region DLK1-DIO3 that contains three clusters with 51 other miRNAs (**Fig. 2A and 2B**). A basewise evolutionary comparison showed miR-2392 is highly conserved among non-human primates such as dogs, cats, and ferrets, species known to be infected with SARS-CoV-2 while mice and rats, species not impacted by COVID-19 (Johansen et al., 2020), have poor conservation with miR-2392.

The impact of miR-2392’s host gene, MEG3, within normal tissues was determined utilizing GTEx data (Consortium, 2020). For the majority of healthy tissues, MEG3 was either not detected or being expressed at low levels (**Fig. 2B**). This can imply that miR-2392 does not seem to significantly affect normal tissues.

We used miRanda (Enright et al., 2003) to identify potential miR-2392 binding sites with respect to the SARS-CoV-2 reference genome (Wuhan-Hu-1; NC045512.2) and lineages of concern. The miR-2392 seeding region is heavily integrated and conserved in different viral strains (**Fig. 2C**). The three best scores are located in the NSP2, NSP3, and E-genes. Notably, these regions were conserved among 6 concerning lineages represented by 14 recent genomes available from the Global Initiative on Sharing All Influenza Data (GISAID, (Shu and McCauley, 2017)).

### MiR-2392 targets mitochondrial and inflammatory pathways associated with SARS-CoV-2

To determine a more comprehensive impact of miR-2392 affected pathways in COVID-19 patients, gene targets were predicted by seed-region base pairing in the miRmap database (Vejnar and Zdobnov, 2012). This list was refined by overlap in several miRNA databases including miRmap, miRwalk (Dweep and Gretz, 2015), miRDB (Chen and Wang, 2020), miRnet (Chang et al., 2020), and ClueGo (Bindea et al., 2009). We added RNA-seq analysis of 39 autopsy tissue samples from the heart, lung, kidney, liver, and lymph node of COVID-19-positive patients with high or low viral loads (Park et al., 2021). MiR-2392 gene targets (375 genes) were visualized using volcano plots (**Fig. 3A-F**).

**Figure 3.**
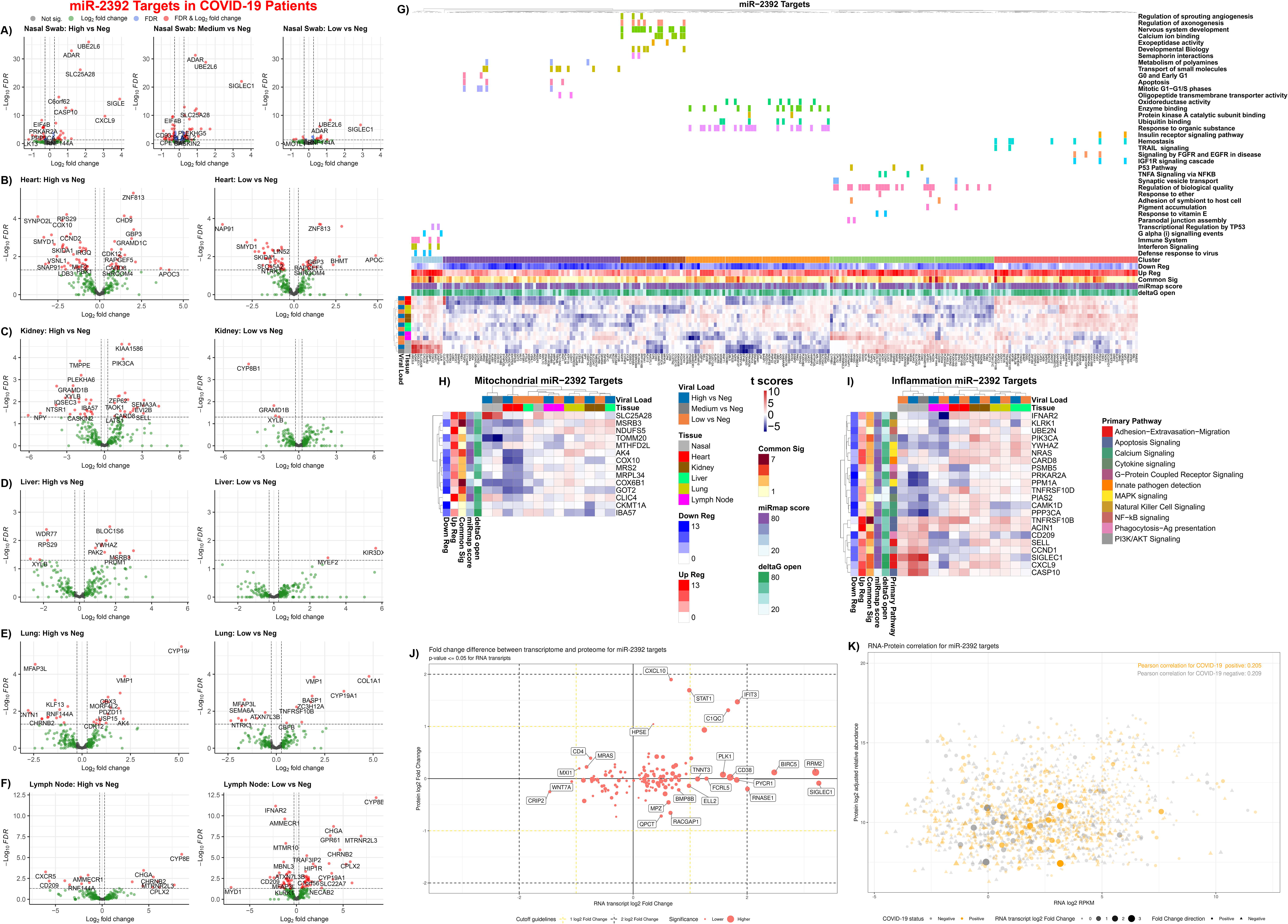
Gene targets of miR-2392 in COVID-19 patients as well as mitochondrial and inflammatory genes. Volcano plots for differential gene expression from **A)** nasopharyngeal swab and autopsy COVID-19 patient tissues from the **B)** heart, **C)** kidney, **D)** liver, **E)** lung, and **F)** lymph node separated by viral load. **G)** Heatmaps display the t-score statistics when comparing viral load versus negative patient samples for nasopharyngeal swab and autopsy tissues. Main gene clusters were determined through k-mean clustering and annotated using ShinyGO (Ge et al., 2020) above the heatmap. MiR-2392 gene targets for **H)** mitochondrial specific genes or **I)** inflammatory genes are displayed. Genes shown have at least one comparison with an adjusted p-value (FDR<0.05) when comparing COVID-19 patients (high, medium or low viral loads) to non-infected control patients (none). Heatmaps from targets determined only from miRmap for miR-2392 mitochondrial or inflammatory gene targets are available in **Fig. S1**. or **Fig. S2**, respectively. **J)** Scatter plot of log_2_-transformed fold changes in RNA and protein for miR-2392 targets for genes differentially expressed at the RNA level. Student’s t-test, RNA p-value<=0.05, no limitation on protein p-value. **K)** Scatter plot of log_2_-transformed medians in RNA and protein. COVID-19 positive (orange) or negative samples (grey). Student’s t-test is used in fold change calculations. Size and opacity of the shape represent RNA fold change values. Shape represents fold change direction: circle = positive, triangle = negative. Pearson correlations are displayed in the top corner.

To further ascertain the systemic impact on miR-2392 gene targets in COVID-19, we performed pathway analysis from the nasopharyngeal swab samples taken from living donors with and without COVID-19 using the SARS-CoV-2 viral load as the independent variable (high, medium, low, other virus). The differentially expressed genes (FDR<0.05) in at least one comparison of COVID-19-positive patients or other detected virus and were found to separate into six distinct hierarchical clusters that were annotated using ShinyGO (Ge et al., 2020) (**Fig. 3G**). The majority of the upregulated miR-2392 targets participate in immune and inflammatory pathways. Downregulated targets are involved in mitochondrial function, oxidative stress, cell cycle, developmental biology, and ubiquitin binding, all pathways recently associated with SARS-CoV-2 infection (Hemmat et al., 2021). This data demonstrates miR-2392 may target several pathways related to perpetuating SARS-CoV-2 infection. For all tissues excluding the lymph nodes, higher viral loads are associated with greater differential expression of miR-2392 regulated gene targets.

Since miR-2392 was recently shown to directly target the transcription of mitochondrial DNA genes (Fan et al., 2019), we evaluated mitochondrial miR-2392 targets in our datasets using MitoCarta (Rath et al., 2021) (**Fig. 3H**). This revealed 14 genes harboring miR-2392 seed sequences that were significantly dysregulated in the nasal and heart samples. In nasal samples, SLC25A28, a mitoferrin which mediates mitochondrial iron transport, was strongly upregulated along with IBA57, which is involved in iron sulfur assembly. The mitochondrial outer membrane protein import complex subunit TOMM20, cytochrome c oxidase (complex IV) subunit COX6B1, and mitochondrial transcription factor COT-2 (NR2F2) were strongly downregulated. In the heart, the folate enzyme MTHFD2L (methylenetetrahydrofolate dehydrogenase) was upregulated while all of the other nuclear-coded mitochondrial genes were downregulated. Downregulated heart mitochondrial genes included NDUFS5 (complex I subunit), COX6B1 and COX10 (complex IV structural and assembly subunits), CKMT1A (mitochondrial creatine kinase), MRPL34 (mitochondrial ribosome small subunit), COT-2 (NR2F2), AK4 and MSRB3 (adenylate kinase 4 and methionine-R-sulfoxide reductase which mitigate oxidative stress), MRS2 (magnesium transporter) and CLIC4 (chloride channel). The kidney showed mild upregulation of complex I and single methyl group metabolism, but down regulation of complex IV (COX10), regulatory factor (COT-2), and iron sulfur center protein (IBA57). Hence, SARS-CoV-2 seems to downregulate nuclear mitochondrial gene transcription in the more oxidative organs, heart and kidney, as well as in nasal tissues.

Since inflammation is a key component of COVID-19, we overlaid known inflammatory genes determined from Loza et al. (Loza et al., 2007) with miR-2392 targets (**Fig. 3I**). From analysis at the mRNA level, most of the complement pathway genes are upregulated in the tissue samples analyzed. These changes could be compensatory, as proteins encoded by the genes could be downregulated as a function of traditional miRNA effects. The responses reflect the importance of degrees of inflammation for mediation of disease severity in COVID-19 patients and a key modulatory role of miR-2392 in this context.

Proteomic and transcriptomic analysis in blood from COVID-19 patients utilizing COVIDome (Sullivan et al., 2021) revealed interesting patterns between RNA and protein levels for miR-2392 targets from miRmap, ClueGO, miRwalk, miRnet, and miRDB (**Fig. 3J and 3K**). Several miR-2392 targets in tissue show a significant transcriptional increase in COVID-19-positive samples with small to no changes on the proteomics level: PLK1, CD38, PYCR1, RNASE1, BIRC5, RRM2, SIGLEC1 (**Fig. 3J**). Interestingly, all these genes were also positively regulated for the majority of tissues when considering only miR-2392 gene targets with miRmap (**Figs. S1 and S2**). The miR-2392 targets CXCL10, STAT1, IFIT3, and C1QC were positively regulated at both the protein and gene levels for the blood and other tissues. The correlation between RNA and protein expression was very close for miR-2392 targets and was slightly stronger in COVID-19 negative (cor=0.209, p=4e-10) versus positive (cor=0.205, p=8e-10) samples (**Fig. 3K**). Further investigation is needed to understand if increased levels of miR-2392 could potentially bind genes’ mRNAs at a higher rate and therefore prevent translation to protein or if there are other mechanisms preventing mRNA translation to protein.

### Overexpression of miR-2392 simulates a phenotype similar to COVID-19 infection

To determine if miR-2392 upregulation would elicit effects similar to a COVID-19 infection, cells were treated with a miR-2392 mimic. Using RNA-seq, 649 genes had a fold-change greater than ±1.2 and a p-value less than 0.05 (**Fig. 4A**) and many genes were miR-2392 predicted targets (**Fig. 4B**). These genes were then compared to whole cell proteome data from a human-derived cell culture model of a SARS-CoV-2 infection profile (Stukalov et al., 2021) and found 10 overlapping gene/proteins that were significantly altered: KIF22, FKBP14, RAD51, AFAP1, ZCCHC17, ZWINT, MAGED1, CENPF, TMEM70, and NFKB2 (**Fig. 4C**). Because viral infection may alter post-translational modification including ubiquitination, we analyzed the ubiquitinome of a SARS-CoV-2 human-derived cell culture model and observed a number of altered proteins in normalized ubiquitin abundance that were also dysregulated genes by miR-2392 overexpression. We also found miR-2392 overexpression impacted genes involved with mitochondria and inflammation (**Fig. 4D-4F**).

**Figure 4.**
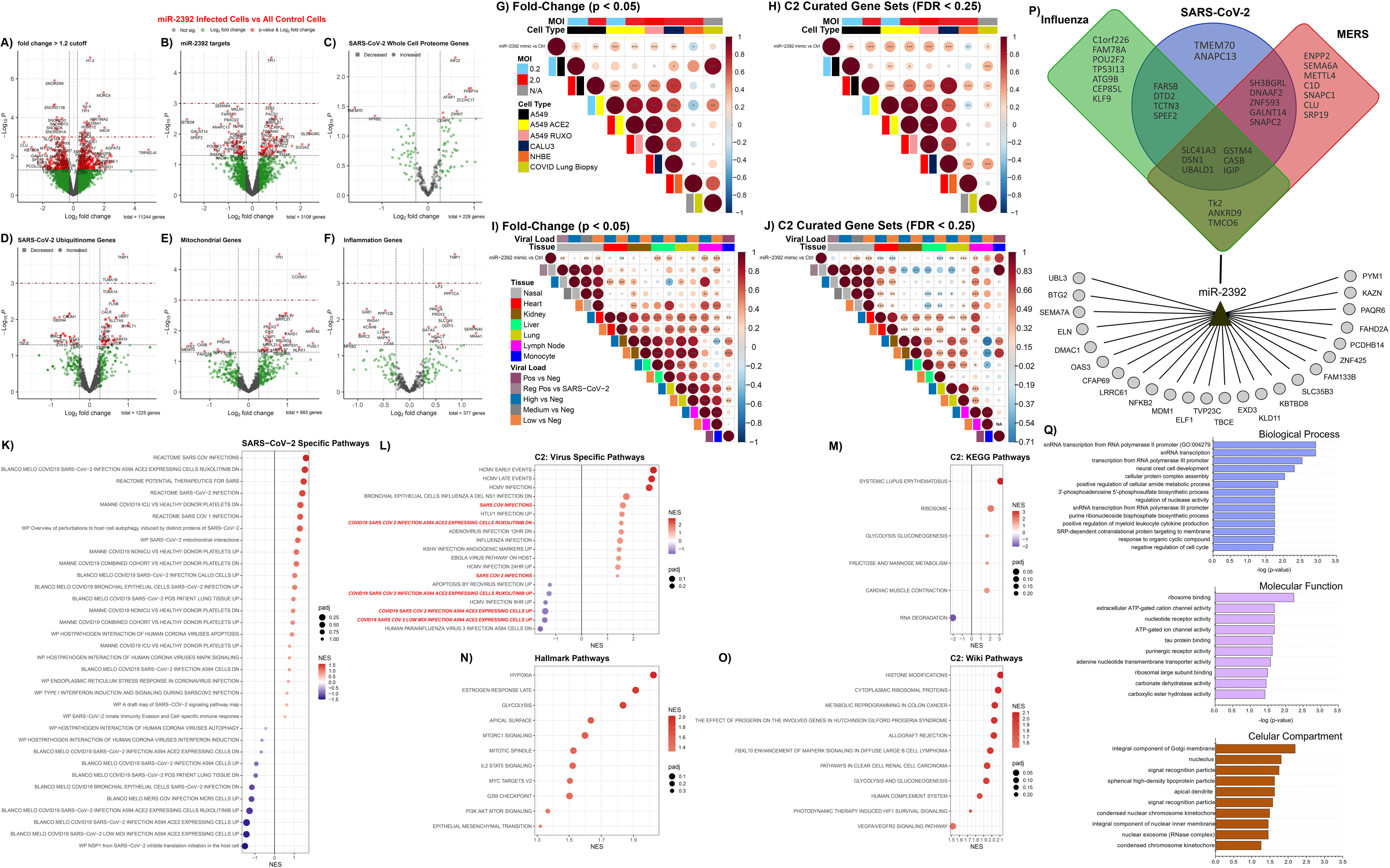
Increased miR-2392 expression *in vitro* mimics a COVID-19 disease phenotype. **A-F)** RNA-seq volcano plots in cells overexpressing miR-2392. **G-J)** Correlation plot between miR-2392 overexpression and related SARS-CoV-2 datasets. Circle size is proportional to the correlation coefficient. Statistical significance determined using a two-tailed Student’s t-test *p<0.05, **p<0.01, ***p<0.001. **K-O**) Dot plots for statistically significant gene sets determined by fGSEA. NES, nominal enrichment score. **P)** and **Q)** Predicted miR-2392 targets by the MIRDIP algorithm that are downregulated in the overexpression experiments. The putative miR-2392 mRNA targets belonging to the consensus transcriptomic networks observed in SARS-CoV-2, MERS and Influenza infections of different human cells are represented in a Venn diagram in the upper part of the panel **P**.

To determine if there was a direct correlation between miR-2392 overexpression and SARS-CoV-2 infection, we compared gene expression changes or overlap in statically significant canonical gene sets with our fGSEA analysis. Published data from Blanco-Melo *et al*. (Blanco-Melo et al., 2020), showed a statistically significant and positive correlation of the miR-2392 treatment to the A549 and Calu-3 cells infected with SARS-CoV-2 (**Fig. 4G and 4H**) as well as in lung biopsies post-mortem from two COVID-19 positive patients (**Fig. 4H**). Using nasal swab samples, there was a significant and positive correlation between patients with medium- and low-viral loads compared to non-infected patients (**Fig. 4I and 4J**). Additional miR-2392 correlation to SARS-CoV-2 infections used RNA-seq data from multiple tissues (heart, kidney, liver, lymph node, and lung) obtained during autopsies of COVID-19 patients with high or low viral loads (**Fig. 4I-J**). There was a positive correlation to lung and lymph node tissues with miR-2392 expression. Interestingly, there was a significant and positive correlation to liver tissue when comparing gene fold-change values (**Fig. 4I**) but not C2 curated biological genesets (**Fig. 4J**). In contrast, a negative correlation to heart tissue was observed.

Statistically significant pathways that were enriched due to miR-2392 treatment were examined using fGSEA **(Fig. 4K-O)**. MiR-2392 treatment induced a pathway response that was significantly related to SARS-CoV-2 pathways. One obvious relationship shows that the Reactome SARS-CoV-2 pathways were significantly activated for the miR-2392 treated cells compared to the controls (**Fig. 4K and 4L**). Significant Hallmark pathways (**Fig. 4N**) show upregulation of hypoxia (Herrmann et al., 2020), glycolysis (Ardestani and Azizi, 2021), and cell cycle pathways (Su et al., 2020) that have been reported to be associated with COVID-19. Interestingly, the KEGG pathway analysis (**Fig. 4M**) indicates miR-2392 overexpression highly upregulated systemic lupus erythematosus which has been reported to occur in COVID-19 patients and have shown similar pathologies due to the increase of inflammation (Zamani et al., 2021).

Lastly, we determined downregulated targets in the cell lines after miR-2392 overexpression. A regulatory network was built by including the predicted miR-2392 targets in the microRNA Data Integration Portal (MIRDIP) that were also downregulated in the overexpression cell model as well as from the recently described consensus transcriptional regulatory networks in coronavirus infected cells (Ochsner et al., 2020) (**Fig. 4P**). The gene enrichment analysis of these putative miR-2392 targets showed the presence of GO-terms related with the RNA metabolism, transcription, ribosome activity and Golgi complex (**Fig. 4Q**).

### Circulating miR-2392 and the suppression of other miRNAs in COVID-19 infected patients

To demonstrate the presence of circulating miR-2392 in COVID-19 infected patients, we quantified the amount of miR-2392 by droplet digital PCR (ddPCR) in the serum, urine and nasopharyngeal swab samples (**Fig. 5**). Serum was obtained from 20 COVID-19 positive patients (10 intubated, 10 not intubated) and ten negative patients. Urine samples were collected from 30 COVID-19 positive patients (15 inpatient, 15 outpatient), 10 inpatient COVID-19 negative samples, and 11 COVID-19 negative healthy donors. Nasal swab samples were obtained from 10 COVID-19 positive patients, 6 common cold coronavirus positive patients (229E, HKU1, and OC43), and 6 Respiratory Illness/Coronavirus NL63 positive patients.

**Figure 5.**
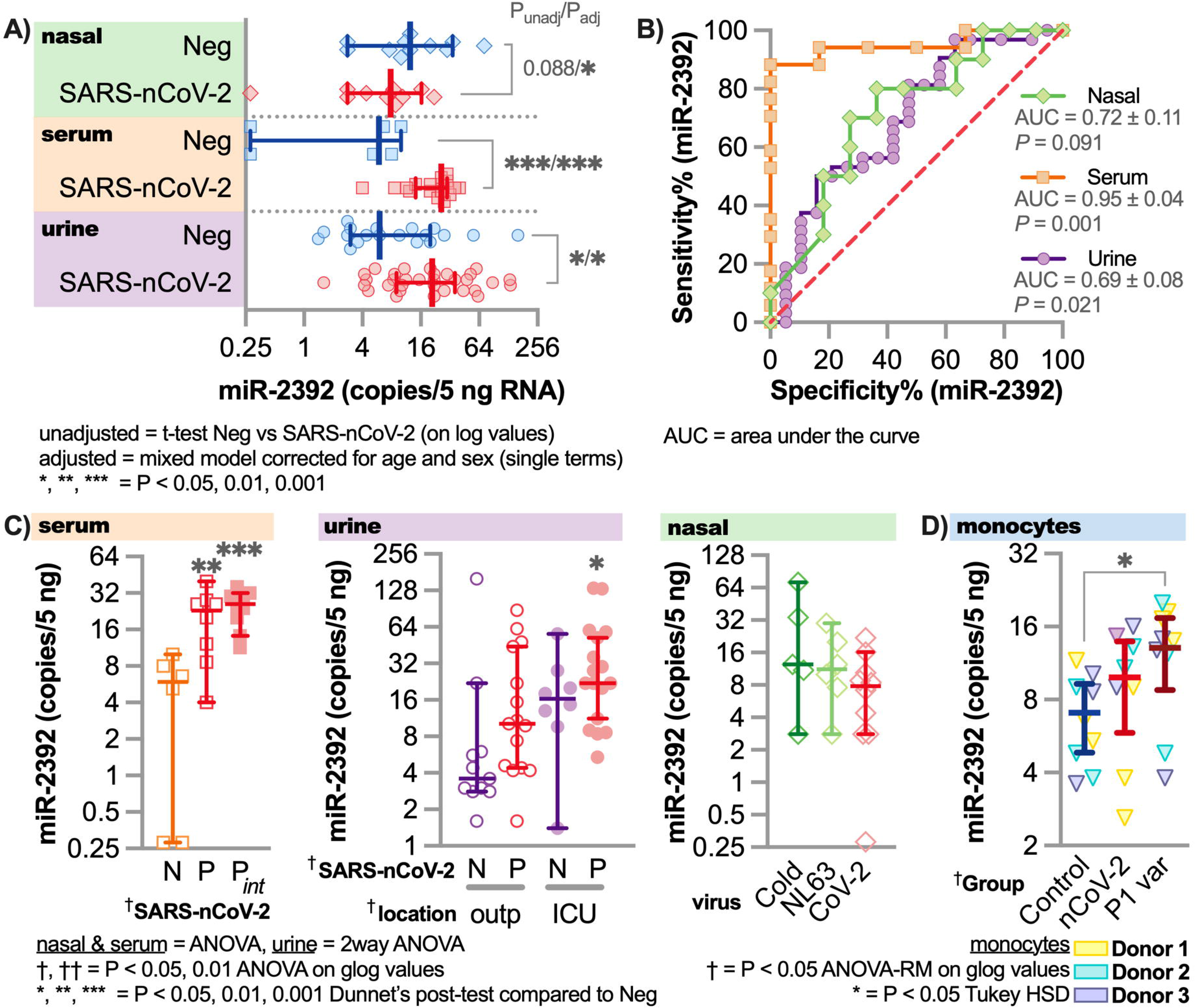
Circulating miR-2392 with COVID-19 patients compared to COVID-19 negative patients. Droplet digital PCR (ddPCR) for miR-2392 on serum, urine, and nasopharyngeal swab samples (including seasonal coronavirus) from COVID-19 positive and negative patients. **A)** MiR-2392 levels from positive (SARS-CoV-2) or negative (neg) patients. Unadjusted t-tests for each tissue or adjusted statistics with a mixed model corrected for age and sex are shown. **B)** ROC curve for miR-2392 in each tissue. **C)** Tissue comparisons for multiple categories. N = COVID-19 Negative, P = COVID-19 positive, P_int_ = intubated patients, outp = outpatient, ICU = Intensive care unit/inpatient, Cold = common cold coronaviruses, NL63 = NL63 coronavirus and CoV-2 = SARS-CoV-2. For all plots *p<0.05, **p<0.01, and ***p<0.001. **D)** MiR-2392 in monocytes from healthy donors (n=3) infected with SARS-CoV-2 reference strain and P1 variant. Comparisons with three other miRNAs were made with miR-1-3p (**Fig. S3**), miR-155-5p (**Fig. S4**), and miR-124-3p (**Fig. S5**).

We observed a statistically significant increase of miR-2392 in COVID-19 positive patients from both the serum and urine samples (**Fig. 5A**). In addition, Receiver Operating Characteristic (ROC) curve analysis revealed that miR-2392 is significantly associated with SARS-CoV-2 infection in patients (**Fig. 5B**) in all tissues. MiR-2392 levels were higher for conditions associated with infection of more severely affected patients (i.e. intubated or ICU) (**Fig. 5C**). Interestingly, low levels miR-2392 appeared in the nasopharyngeal location with no significant differences occurring between seasonal coronavirus samples. To study more specific tissue origins for miR-2392 we infected monocytes obtained from healthy donors with SARS-CoV-2 reference strain (i.e. strain from Wuhan) and also P1 variant (**Fig. 5D**). We saw an increase of miR-2392 with the reference SARS-CoV-2 infected cells and a significant increase in the P1 variant infected cells. Previously, it was shown that these monocytes infected with SARS-CoV-2 have increased pro-inflammatory cytokine and glycolysis expression, with the effects inhibited by glycolysis inhibitors (Codo et al., 2020). Since we hypothesize that miR-2392 is a primary initiator for systemic impact of the infection, this might indicate that miR-2392 does not strongly appear until the virus has established its presence in the body.

We also quantified miR-1-3p (**Fig. S3**), miR-155-5p (**Fig. S4**), and miR-124-3p (**Fig. S5**) which were predicted to be inhibited by COVID-19 infection (**Fig. 1A**). MiR-1-3p and miR-155-5p were significantly suppressed in the serum with no differences in the urine or nasopharyngeal samples (**Fig. S3 and S4**). MiR-1-3p is known to be beneficial for cardiovascular functions, with its inhibition leading to heart failure and disease (Condorelli et al., 2010). Interestingly, when quantifying miR-1-3p in the SARS-CoV-2 infected monocytes we observe a significant decrease with the reference strain, while no different with the P1 variant (**Fig. S3D**). For miR-124-3p, we observed very low amounts (on average < 2 copies/5 ng RNA), for all conditions, which indicates that miR-124-3p is not circulating for any of the patients for any the conditions observed (**Fig. S5**). MiR-124-3p provides an ideal miRNA negative control candidate for SARS-COV-2.

### Inhibiting miR-2392: a novel antiviral COVID-19 therapeutic

The link between miR-2392 and COVID-19 infection prompted the develop of an effective antiviral approach for COVID-19 by inhibiting miR-2392. We used the Nanoligomer platform to develop an effective antisense-based therapeutic against human miR-2392 (Eller et al., 2021), termed SBCov207 (**Fig. 6A**). The anti-miR-2392 Nanoligomers was evaluated for efficacy and toxicity against a SARS-CoV-2 infection of the human lung cell line A549 (**Fig. 6B-D**). Treatment of uninfected A549 cells showed no cytotoxicity up to 20 µM. The control nonsense Nanoligomers (SBCoV208) showed no toxicity even up to 40 µM. Treatment of A549 cells infected with SARS-CoV2 showed drastic improvement in cell viability with an average of 85% viral inhibition at 10 µM (IC50=1.15±0.33 µM). In contrast, the control Nanoligomers showed significantly lower viral suppression (**Fig. 6E-G**). This human cell line model reaffirms that the anti-miR-2392 (SBCov207) is effective in inhibiting SARS-CoV2, while not exhibiting toxicity at the concentrations tested.

**Figure 6.**
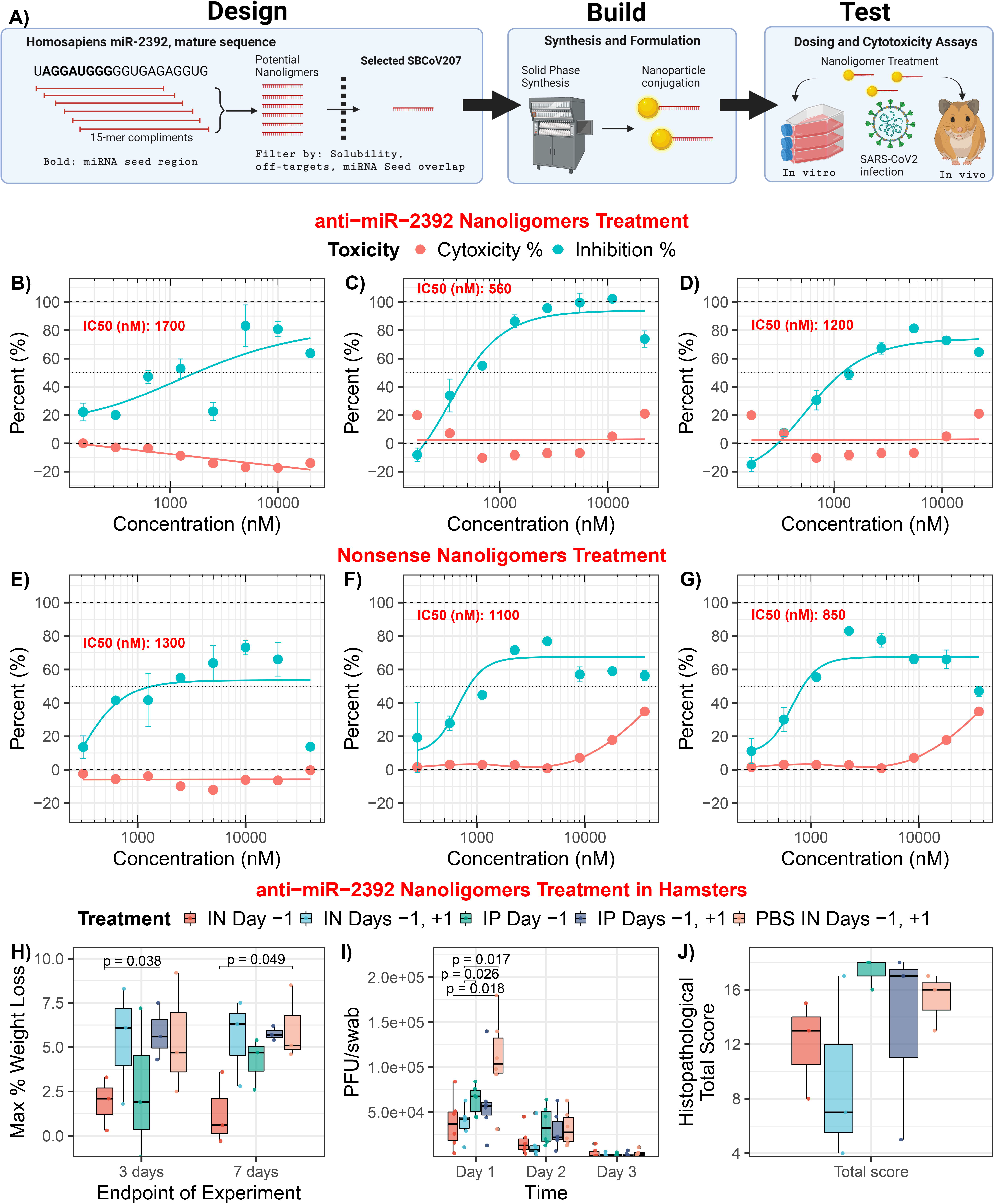
Anti-miR-2393 therapeutic mitigation of SARS-CoV-2 infection with *in vitro* and *in vivo* models. **A)** Platform schematic for the miR-2392 inhibitor. **B–G)** Anti-miR-2392 Nanoligomers inhibitor or **E–G)** nonsense control applied to A549 cells infected with SARS-CoV-2 and tested for viral viability and cytotoxicity as biological replicates (n=3). **H–J)** Toxicity and efficacy of anti-miR-2392 Nanoligomers inhibitor in an *in vivo* infection hamster model. Treatments groups: SBCov207 by IP injection or IN instillation 24 hours prior to viral inoculation (IP or IN Day -1), 24 hours prior and post-viral challenge (IP or IN Day -1, +1), and 100ul of PBS 24 hours prior and post-viral challenge through IN instillation (PBS IN Day -1, +1). **H)** Pooled weights for each treatment group (n = 6 for days 1 – 3 and n = 3 for days 4 -7) as the maximum percent weight loss. **I)** SARS-CoV-2 assayed by plaque assay on Vero E6 cells from oropharyngeal swabs (N=6). **J)** Histopathological total score for lung tissues at day 3. Intranasal (IN), intraperitoneal (IP).

The anti-miR-2392 Nanoligomers was then evaluated in a Syrian hamster infection model (**Fig. 6H-6J**). Six hamsters were treated with Nanoligomers for 72 hours without infection and there was no observed change in animal behavior indicating a lack of obvious toxicity. Next, 30 male hamsters were divided into 5 treatment groups. The infected hamsters were given 10^5^ plaque forming units (pfu) of WA01/2020 strain of SARS-CoV-2. The anti-miR-2392 Nanoligomers treatment was given by intraperitoneal (IP) injection or intranasal (IN) instillation once at 24 hours before or twice at 24 hours before and after viral inoculation. Half of the hamsters in each group (n=3) were euthanized and necropsied on day 3 and 7 post-infection respectively.

Loss of body weight over the course of the experiment was <10% in all groups and significantly different for the IN treatment one day before viral inoculation (compared to the control) while statistical differences in other groups were not present (**Fig. 6H**). Virus titers from oropharyngeal swabs for IN treatment were significantly lower (*p* = 0.018) than those from hamsters receiving Nanoligomers IP or PBS on day 1 post-challenge, but there were no differences among groups on days 2 and 3 post-challenge (**Fig. 6I**). Although not statistically different than the control treatment, the data indicates a downward trend with Nanoligomers treatment (**Fig. 6J**) and the total histopathological score for the IN was lower than the controls.

### The impact of miR-2392 on diseases, relationship to COVID-19 symptoms, and predicted FDA drugs to target miR-2392

To predict whether miR-2392 might have a direct relationship to COVID-19 symptoms, we determined the pathway and disease relevance using miRnet. Among the diseases predicted to be associated with miR-2392 were a surprising number of clinical observations present in individuals with COVID-19 infection (**Fig. 7A**). These include heart or cardiovascular disease and failure, both known to heavily contribute to morbidity and mortality in patients with COVID-19 (Nishiga et al., 2020), hyperesthesia (Krajewski et al., 2021), as well as less common COVID-19 symptoms, such as lymphadenopathy and pharyngitis related to sore throat (Edmonds et al., 2021), liver dysfunction (Portincasa et al., 2020), splenomegaly (Malik et al., 2020), CNS (Mahajan and Mason, 2021; Rodriguez et al., 2020) and kidney failure (Hultstrom et al., 2021).

**Figure 7.**
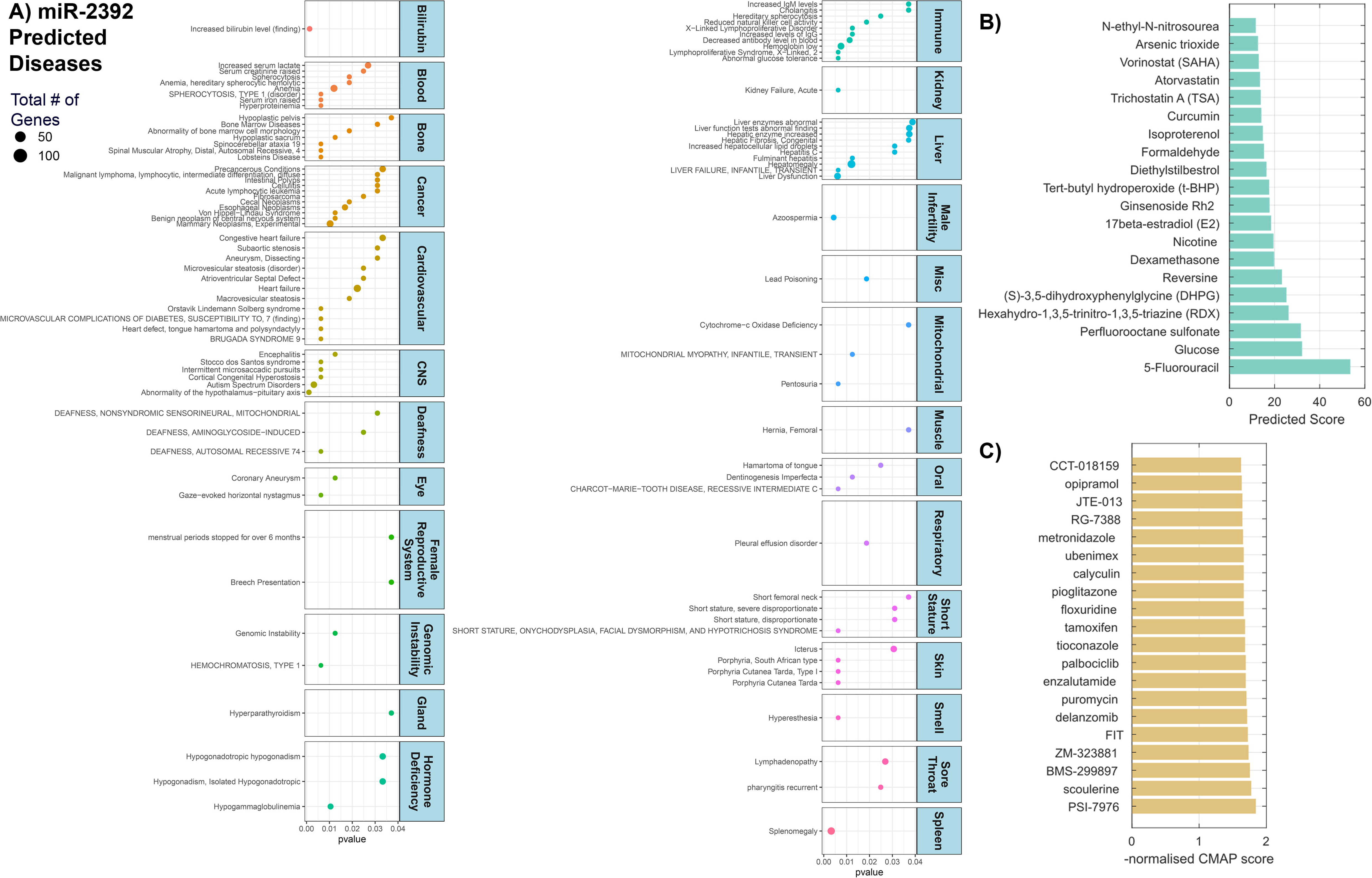
Predicted impact of miR-2392 on human disease and the top-20 drug compounds predicted to affect miR-2392 expression through machine learning approach. **A)** Dot plot of manually curated diseases associated with miR-2392 predicted from miR-2392 gene targets by miRnet. Values are plotted by p-value and dot size represents the number of downstream gene targets associated with disease. MiR-2392 expression related to patient survival in a pan-cancer analysis is highlighted in **Fig. S7**. **B)** Barplot of scores using our matrix completion model to predict small molecules that affect miRNA expression. Higher scores indicate more predicted associations. **C)** Barplot of the normalized connectivity map (CMAP) scores. We used transcripts induced by miR-2392 overrepresented genes to query CMAP. Higher negative scores reflect a greater reversal of the miR-2392 transcriptomic signature. Further details on model statistics and performance are found in **Figs. S8 – S10**.

MiR-2392 was also predicted to affect other diseases that have been linked to COVID-19 infection in some patients. For example, COVID-19 patients have experienced azoospermia, which is linked to male infertility (Younis et al., 2020), altered menstrual cycle in females (Li et al., 2021), dental damage (Sirin and Ozcelik, 2021), and deafness or hearing loss (Koumpa et al., 2020). Using the tool Kaplan-Meier Plotter (Győrffy, 2021) to associate miR-2392 expression with pan-cancer patient survival (**Fig. S7**), a high expression of miR-2392 was generally related to poor prognosis with the majority of cancer types (p<0.05). Intriguingly, one miR-2392 predicted consequence was decreased antibody levels in the blood; this might account for the reported loss of the antibodies overtime (Gudbjartsson et al., 2020; Self et al., 2020).

Finally, using computational models we predicted small molecules, including FDA-approved drugs, that could inhibit miR-2392. First, a state-of-the-art machine learning algorithm for predicting missing drug targets (Galeano et al., 2021) was applied to an association dataset between 213 small molecules and 1,519 miRNAs from the SM2miR database (Liu et al., 2013) (see statistics in **Fig. S8**) integrating chemical and sequence similarity respectively. The average area under the ROC was of 0.877 when predicting missing small molecule-miRNA associations (**Fig. S9**). The top-20 predicted small molecules (**Fig. 7B**) includes Dexamethasone (Ledford, 2020) and Atorvastatin, that have shown protective roles in COVID-19 patients (Rossi et al., 2020). Second, following the ideas first presented in Sirota *et al*. (Sirota et al., 2011), we performed an analysis on the genomic signature of miR-2392 (*i.e.* significant up and down-regulated genes) and predicted small molecules that can reverse it. We screened the genomic signatures of miR-2392 against 30,000 small molecules contained in the connectivity map (CMAP) (Lamb et al., 2006). The top-20 predicted small molecules includes the androgen receptor antagonist Enzalutamide and the insulin sensitizer Pioglitazone (Carboni et al., 2020) both of which are in clinical trials for COVID-19 (**Fig. 7C;** Clinical Trial NCT04475601 and NCT04604223). We also found literature evidence for leukotriene inhibitor ubenimex (Asai et al., 2020), and bacterial DNA inhibitor metronidazole (Gharebaghi et al., 2020).

## Discussion

While the potential eradication of the novel coronavirus through worldwide vaccination is underway, new potent strains of SARS-CoV-2 are constantly evolving and there remains a major need to develop effective interventional strategies to minimize the damage caused by coronavirus infections. Host-mediated lung inflammation is a driver of mortality in COVID-19 critically ill patients. Thus, it is logical to focus on therapeutics that may have immunomodulating properties or disrupt viral replication. Our research uncovers a novel eight miRNA signature in patients with COVID-19 viral loads compared to those without disease as predicted from RNA-seq data. The expression of seven miRNAs was decreased (miR-10, miR-1, miR-34a-5p, miR-30c-5p, miR-29b-3p, miR-124-3p, and miR-155-5p) while a single miRNA, miR-2392, was significantly increased (**Fig. 1**). This key miRNA signature was involved in major cellular and molecular mechanisms that drives the viral-host response.

Upregulation of miR-10a-5p led to suppression of SDC1 defending against Porcine hemagglutinating encephalomyelitis virus (Hu et al., 2020). Notably, upregulation of mir-30c-5p and miR-155-5p have been independently shown to be involved with antiviral functions through immune and inflammatory pathways with other coronaviruses (Dickey et al., 2016; Ma et al., 2018). Inhibition of miR-34a-5p in the host by SARS-CoV-2 suppresses beneficial antiviral pathways (Bartoszewski et al., 2020). MiR-1-3p has been identified as an antiviral agent for viral related respiratory diseases (Sardar et al., 2020). MiR-1-3p and miR-155-5p from serum samples were significantly inhibited (**Figs. S3 and S4**), in agreement with the literature above.

Several studies have measured differential expression of miRNAs in COVID-19 patients and proposed their use as biomarkers or therapeutics. Lung biopsies from 9 COVID-19 patients showed miR-26a, miR-29b, and miR-34a correlated to endothelial dysfunction and inflammatory biomarkers (Centa et al., 2020). Sequencing in the blood from moderate or severe COVID-19 patients identified miR-146a, miR-21, miR-142, and miR-15b as potential biomarkers as well as contributors to disease pathogenesis (Tang et al., 2020). While these studies are limited to a specific tissue, our data that correlates miRNA signatures from multiple tissues (**Fig. 3**) suggests miR-2392 is a unique target that is ubiquitously involved in COVID-19 symptoms. To start addressing the multi-tissue impact of miR-2392 we show that miR-2392 is overexpressed in healthy monocyte cells infected with both the reference and P1 SARS-CoV-2 variants (**Fig. 5D**). This can indicate isolated cells will overexpress miR-2392 when infected with SARS-CoV-2 and further work is currently being done to investigate multi-tissue behavior of miR-2392.

The majority of miR-2392 publications are focused on cancer tissues where it may drive cellular invasion and metastasis. MiR-2392 was one of 6 circulating miRNAs altered in the serum and tissue of patients with cervical cancer used to predict metastasis (Chen et al., 2013). Higher levels of miR-2392 in gastric cancer was associated with lower clinical staging and increased patient survival (Li et al., 2017) by inhibiting invasion and metastasis through the loss of Snail1, Slug, and Twist1 expression. Similarly, lower miR-2392 and miR-1587 levels were found in human keloid tissues that resulted in a loss of inhibition of ZEB2 and subsequent promotion of cellular proliferation and invasion (Hou et al., 2017). Inhibition of miR-2392 by the long-non-coding RNA CACNA1G-AS1 promoted hepatocellular carcinoma through disrupting the degradation of C1orf61, a tumor activator associated with metastasis and progression (Hu et al., 2013; Yang et al., 2019). Recently, a novel role for miR-2392 in tongue squamous cell carcinoma showed partial inhibition of transcription through direct miRNA-mtDNA base pairing resulting in reprogramming tumor cell metabolism and chemoresistance (Fan et al., 2019). These miR-2392 reports establish the significant impact this miRNA may have on cellular activity. Particularly relevant here is the more than 2-fold increase of miR-2392 in extracellular vesicles of Hepatitis B infected human hepatocytes (Enomoto et al., 2017). While miR-2392 has a reported impact on tumor cell biology, our study expands the valuable therapeutic potential of targeting miR-2392 to subsequently decrease SARS-CoV-2 viral infections (**Fig. 6**). These results warrant further exploration of the mechanistic underpinnings for the role of miR-2392 in driving viral infection.

One therapeutic insight deduced from miR-2392 interactions is the importance of the mitochondrial oxidative phosphorylation (OXPHOS) and glycolytic pathways in COVID-19, dramatically highlighted in BALF samples (**Fig. 1C**). In a study of tongue squamous cell carcinoma (Fan et al., 2019), miR-2392 enters the mitochondrion where it binds Ago2 and nucleotides 4379 to 4401 in the mtDNA heavy (H) strand within the MT-TQ (tRNA glutamine) gene (nucleotides m.4329-4400). MT-TQ is part of a large polycistronic transcript encompassing 12 of the mtDNA H strand polypeptide genes punctuated by tRNAs. Cleavage of the tRNAs releases the mRNAs. Upstream of MT-TQ are the 12S and 16S rRNAs and the complex I gene MT-ND1 gene. Downstream of MT-TQ is MT-ND2, MT-CO1, MT-CO2, MT-ATP6/8, MT-ND3, MT-ND4L, MT-ND4, MT-ND5, and MT-CYB (Lott et al., 2013; Wallace, 2018). Strikingly, the down-regulated mtDNA genes are the complex IV (cytochrome c oxidase) genes MT-CO1 and MT-CO2, the complex III (the bc_1_ complex) gene (MT-CYB), and the complex I genes (MT-ND2, MT-ND4, and MT-ND5) (**Fig. 1C** right side arc). Since the miR-2392 inhibition of mtDNA OXPHOS genes (**Fig. 1C**) is also reflected in the down-regulation of the nuclear DNA coded mitochondrial transcripts of the complex I and IV genes and the iron-sulfur and heme iron complexes in the nasal, heart, and kidney autopsy samples (**Fig. 3D**), mitochondrial inhibition by miR-2392 appears to be the only physiological function that is common across all tissues in infected individuals suggesting mitochondrial modulation is a central feature of SARS-CoV-2 pathophysiology.

The inhibition of mitochondrial genes by miR-2392 would impair OXPHOS, which would have the most adverse effects on the high mitochondrial energetic tissues (brain, heart, kidney), the tissues central to the most severe COVID-19 cases. Inhibition of mitochondrial OXPHOS genes would increase mitochondrial reactive oxygen species (mROS) production, and induce glycolysis to compensate for the energy deficit (see top of **Fig 1C**). Mitochondrial function is regulated by the Sirtuins (Carrico et al., 2018), mitochondrial decline is associated with senescence, and mROS oxidation of mtDNA is linked to activation of the inflammasome and thus NFκB (West and Shadel, 2017; Zhong et al., 2018), all of which are modulated around miR-2392 (**Fig. 1C**). Thus, SARS-CoV-2 induction of miR-2392 (**Fig. 5**) and its associated inhibition of mtDNA and nuclear DNA OXPHOS genes (**Fig. 3** and **S1**) could explain many of the metabolic disturbances of COVID-19. Conversely, antagonism of miR-2392 function should ameliorate the inhibition of OXPHOS and may explain the therapeutic benefit of the anti-miR-2392 Nanoligomers.

Exploiting miRNAs from serum as a biomarker was pioneered in patients to examine diffuse large B-cell lymphoma (Lawrie et al., 2008). The use of miRNAs as a diagnostic biomarker has several advantages. Circulating miRNAs are readily obtained through a minimally invasive blood draw and are remarkably resistant to degradation in the plasma and serum (Mitchell et al., 2008). They may provide a means to detect asymptomatic individuals (Hou et al., 2017). However, potential confounding diseases that may influence the expression of multiple miRNAs requires the further evaluation of the targets found in this study (**Fig. 5**).

Advances in RNA chemistry and delivery systems enabled the first miRNA-based agents to enter into clinical trials several years ago (Rupaimoole and Slack, 2017). MiR-122 was found to increase the stability and replication of the Hepatitis C virus (HCV) through binding that prevented degradation by the Xrn1 exoribonuclease (Jopling et al., 2005; Thibault et al., 2015). Following a phase I clinical trial that inhibited miR-122 with no adverse reactions, a phase IIa trial showed a significant dose-dependent decrease in HCV load, one reported grade 3 thrombocytopenia, and only a small set of patients experienced viral rebound that might be linked to mutations of the HCV viral RNA (Janssen et al., 2013; Ottosen et al., 2015). A separate clinical trial targeting miR-122 reduced viral loads in all treated patients within 4 weeks with a sustained response in 3 patients after 76 weeks (van der Ree et al., 2017), however trials were suspended due to two cases of severe jaundice. These clinical trials have demonstrated the promising potential of using anti-miRNAs to significantly reduce viral infection with limited adverse effects and the similarities with miR-2392 with SARS-CoV-2 warrant further investigations to push to clinical trials.

Presently, there remains no specific treatment option for patients presenting with severe COVID-19 disease. While vaccines provide a promising avenue towards curbing the infection rate and minimizing symptoms, there remains an urgency to successfully develop and implement therapeutic agents to reduce severe consequences and subsequent patient mortality from COVID-19. As the testing of antibody-based or drug targeted therapies are currently underway, miRNAs represents a novel category of therapeutic agents that have previously shown endogenous activity to alter viral infection.

## Acknowledgments

This work was supported by supplemental funds for COVID-19 research from Translational Research Institute of Space Health through NASA Cooperative Agreement NNX16AO69A (T-0404 to A.B. and T-0406 to A.C.) and further funding from KBR, Inc provided to A.B; this work used resources services, and support provided via the COVID-19 HPC Consortium (https://covid19-hpc-consortium.org/), provided specifically by the NASA High-End Computing (HEC) Program through the NASA Advanced Supercomputing (NAS) Division at Ames Research Center which was awarded to A.B.; DOD W81XWH-21-1-0128 awarded to D.C.W.; NIGMS P20 GM1009005 awarded to N.A.; Individual National Research Service Award F32-AI147587 awarded to J.M.D.; NIH/NHBLI K08 HL143271 and NIH/NHLBI R03 HL155249 awarded to R.S.H.; NIH/NCI U54 CA260543 supported R.S.H., N.M.B., M.C.W.; NSF 1956233 awarded to R.M.; A.P. was supported by Biotechnology and Biological Sciences Research Council (https://bbsrc.ukri.org/) grants BB/K004131/1, BB/F00964X/1 and BB/M025047/1, Consejo Nacional de Ciencia y Tecnología Paraguay - CONACyT (http://www.conacyt.gov.py/) grants 14-INV-088 and PINV15-315, National Science Foundation Advances in Bio Informatics (https://www.nsf.gov/) grant 1660648; NSF IOS 1546858 to ESW; D.G. and A.P. were supported by the CONACyT grant PINV20-337 and the Fundação Getulio Vargas. N.M.B. was supported by the National Center for Advancing Translational Sciences (NCATS), NIH UL1TR002489, 2KR1272005, 550KR242003. P.M.M.V. was supported by FAPESP (grant numbers 20/04579-7, 16/18031-8); Fundo de Apoio ao Ensino, Pesquisa e Extensão (FAEPEX), Unicamp (grant number 2274/20); Coordenação de Aperfeiçoamento de Pessoal de Nível Superior - Brazil (CAPES) (Finance Code 001). Thanks to Dr. Richard Bowen for his assistance with the hamster studies and Dr. Matthew Frieman for his assistance with the *in vitro* A549 SARS-CoV-2 studies. Graphical abstract was created with BioRender.com partially adapted from “Bevacizumab: Potential Repurposed Drug Candidate for COVID-19” retrieved from https://app.biorender.com/biorender-templates.

## Author contributions

Conceptualization: A.B.; Methodology: A.B.; Formal Analysis: A.B., R.M., C.V., D.T., F.J.E., C.M., C.M., J.C.S., J.T.M., J.M.D., D.G., U.S., E.S.W., A.S., J.F., V.Z., N.S., T.J.T.; Investigation: A.B., C.V., R.M., C.E.M., A.C., P.N., S.L.M., A.Y., T.R.A., P.M.M.V., G.G.D.; Sample Collection: M.M.S., M.C.W., R.S.H., N.M.B., U.C.P.C., A.D.H., J.C. P.M.M.V., G.G.D.; Resources: A.B., R.M., C.E.M., A.C., M.S., M.F., M.C.W.; Writing – Original Draft: A.B., J.T.M.; Writing – Review & Editing: A.B., J.T.M., F.J.E., R.J.G., W.P., M.M.S., J.M.D., J.W.G., D.W., S.S., S.B., V.Z., E.S.W., S.V.C., N.A, A.P., D.G., P.M.C., M.R.E., J.C.S., A.C., R.M., N.S., T.J.T., B.C., L.N.S., M.C.W., P.M.M.V.; Visualization: A.B., J.S.C, F.J.E, D.G., N.S., V.Z.; Supervision: A.B.; Funding Acquisition: A.B.

## Declaration of Interests

A.C., P.N., S.V.C., A.B. have a provisional patent based on the antiviral discovery and design. A.C., P.N. and S.S. are part of the company (Sachi Bioworks Inc.) that has filed a patent on the Nanoligomer technology.

## Supplemental Figures and Material

**The UNC COVID-19 Pathobiology Consortium is comprised of:**

Shannon M Wallet^1,2^, Robert Maile^2,3^, Matthew C Wolfgang^2,4^, Robert S Hagan^4,5^, Jason R Mock^4,5^, Natalie M Bowman^6^, Jose L Torres-Castillo^5^, Miriya K Love^5^, Suzanne L Meinig^4^, Will Lovell^1^, Colleen Rice^5^, Olivia Mitchem^1^, Dominique Burgess^1^, Jessica Suggs^1^, Jordan Jacobs^3^

^1^Department of Oral and Craniofacial Health Sciences, UNC Adams School of Dentistry, University of North Carolina School of Medicine, Chapel Hill, NC, 27599, USA

^2^Department of Microbiology & Immunology, University of North Carolina School of Medicine, Chapel Hill, NC, 27599, USA

^3^Department of Surgery, University of North Carolina School of Medicine, Chapel Hill, NC, 27599, USA

^4^Marsico Lung Institute, University of North Carolina at Chapel Hill, Chapel Hill, NC, 27599, USA

^5^Division of Pulmonary Diseases and Critical Care Medicine, Department of Medicine, University of North Carolina, Chapel Hill, NC 27599, USA

^6^Division of Infectious Disease, School of Medicine, University of North Carolina, Chapel Hill, NC, 27599, USA

**Figure S1**. Mitochondrial gene targets of miR-2392 and regulated pathways. Related to Figure 3. Differential gene expression analysis for all miR-2392 mitochondrial gene targets significantly expressed in nasopharyngeal swab and autopsy COVID-19 patient tissues. The heatmaps display the t-score statistics for comparing viral load vs negative patient sample for all samples. Main gene clusters were determined through k-mean clustering. Nine main gene clusters were determined and ShinyGO (Ge et al., 2020) was utilized to determine the pathways for each cluster which are displayed on the top panel of the heatmap. Differentially expressed genes are shown with at least one comparison demonstrating a significant adjusted p-value (FDR<0.05) when comparing COVID-19 patients (high, medium or low viral loads) to non-infected control patients (none). Mir-2392 gene targets only determined from miRmap.

**Figure S2**. Inflammatory gene targets of miR-2392 and regulated pathways. Related to Figure 3. Differential gene expression analysis for all miR-2392 inflammatory gene targets significantly expressed in nasopharyngeal swab and autopsy COVID-19 patient tissues. The heatmaps display the t-score statistics for comparing viral load vs negative patient sample for all samples. Main gene clusters were determined through k-mean clustering. Eight main gene clusters were determined and ShinyGO (Ge et al., 2020) was utilized to determine the pathways for each cluster which are displayed on the top panel of the heatmap. Differentially expressed genes are shown with at least one comparison demonstrating a significant adjusted p-value (FDR<0.05) when comparing COVID-19 patients (high, medium or low viral loads) to non-infected control patients (none). Mir-2392 gene targets only determined from miRmap.

**Figure S3**. Circulating miR-1-3p with COVID-19 patients compared to COVID-19 negative patients. Related to Figure 5. Droplet digital PCR (ddPCR) with specific primer for miR-1-3p was performed on serum, urine, and nasopharyngeal swab samples (including other seasonal coronavirus samples) from COVID-19 positive and negative patients. The miRNA concentration are reported as copies/5ng RNA. A) The levels of miRNA-2392 in all tissues from patients grouped as SARS-CoV-2 positive (SARS-nCoV-2) or negative (neg). Unadjusted t-tests comparing the SARS-CoV-2 positive to neg for each tissue are provided and also adjusted statistics comparing the groups with a mixed model corrected for age and sex is provided. B) Receiver Operating Characteristic (ROC) curve is provided for miR-1-3p for each tissue comparing SARS-CoV-2 positive to negative patients. C) Comparing specific categories within each tissue type between COVID-19 positive and negative patients. N = COVID-19 Negative, P = COVID-19 positive, P_int_ = intubated patients, outp = outpatient, ICU = Intensive care unit/inpatient, Cold = Coronaviruses related to the common cold, NL63 = NL63 coronavirus, and CoV-2 = SARS-CoV-2. D) The levels of miR-1-3p in monocytes from healthy donors (n = 3) that were infected with SARS-CoV-2 reference strain and P1 variant. Triplicate conditions were done for each donor. For all plots * = p < 0.05, ** = p < 0.01, and *** = p < 0.001.

**Figure S4**. Circulating miR-155-5p with COVID-19 patients compared to COVID-19 negative patients. Related to Figure 5. Droplet digital PCR (ddPCR) with specific primer for miR-155-5p was performed on serum, urine, and nasopharyngeal swab samples (including other seasonal coronavirus samples) from COVID-19 positive and negative patients. The miRNA concentration are reported as copies/5ng RNA. A) The levels of miRNA-2392 in all tissues from patients grouped as SARS-CoV-2 positive (SARS-nCoV-2) or negative (neg). Unadjusted t-tests comparing the SARS-CoV-2 positive to neg for each tissue are provided and also adjusted statistics comparing the groups with a mixed model corrected for age and sex is provided. B) Receiver Operating Characteristic (ROC) curve is provided for miR-155-5p for each tissue comparing SARS-CoV-2 positive to negative patients. C) Comparing specific categories within each tissue type between COVID-19 positive and negative patients. N = COVID-19 Negative, P = COVID-19 positive, P_int_ = intubated patients, outp = outpatient, ICU = Intensive care unit/inpatient, Cold = Coronaviruses related to the common cold, NL63 = NL63 coronavirus, and CoV-2 = SARS-CoV-2. For all plots * = p < 0.05, ** = p < 0.01, and *** = p < 0.001.

**Figure S5**. Circulating miR-124-3p with COVID-19 patients compared to COVID-19 negative patients. Related to Figure 5. Droplet digital PCR (ddPCR) with specific primer for miR-124-3p was performed on serum, urine, and nasopharyngeal swab samples (including other seasonal coronavirus samples) from COVID-19 positive and negative patients. The miRNA concentration are reported as copies/5ng RNA. For miR-124-3p, the copies/5ng were either equal to 0 or at extremely low levels close to 0 copies/5ng. To try to determine any statistical differences we categorized the groups as ND = Not Determined which are all 0 values or D = Determined which are values > 0 for both N = negative (open symbols) and P = COVID-19 positive patients samples (closed symbols). The number of patients for each column is shown above the points. No significant differences were observed for any of the sample for miR-124-3p.

**Figure S6**. miR-2392 expression pan-cancer survival analysis. Related to Figure 7. Kaplan Meier patient survival plots for miR-2392 expression in a pan-cancer analysis was determined utilizing The Kaplan Meier plotter (Nagy et al., 2021). The plots were separated with the top row being cancers which patients had significantly poor survival with high expression of miR-2392, the middle row being cancers which patients had poor survival (but not significant) with high expression of miR-2392, and the bottom row being cancers which patients had significantly better survival with high expression of miR-2392.

**Figure S7**. Small molecules-miRNA dataset statistics Related to Figure 7. (Left) Number of small molecules associated to miRNAs. (Right) Number of miRNAs associated to small molecules.

**Figure S8**. Performance of our method at predicting missing small molecule-miRNA interactions. Related to Figure 7. (Top) The mean value of the Receiver Operating Curve (ROC) is shown for a ten-fold cross-validation experiment (dark blue). 95% confidence interval is also shown (light blue). (Bottom) The mean value of the Precision-Recall Curve (PRC) is shown for a ten-fold cross-validation experiment (dark salmon). 95% confidence interval is also shown (light salmon).

**Figure S9**. Performance of our method at predicting missing small molecule-miRNA interactions when controlling for data imbalance. Related to Figure 7. (Top) Area Under the Receiver Operating Curve (AUROC) was obtained in a ten-fold cross-validation experiment for varying values of the negative to positive label ratio in the test set. (Bottom) Area Under the Precision-Recall Curve (AUROC) was obtained in a ten-fold cross-validation experiment for varying values of the negative to positive label ratio in the test set.

**Table S1. Annealing temperatures for miRNA primers, related to methods and Figure 5.** Temperatures used for droplet digital PCR to quantify each miRNA target.

## STAR★Methods

### RESOURCE AVAILABILITY

#### Lead Contact

Further information and requests for resources and reagents should be directed to and will be fulfilled by the Lead Contact, Afshin Beheshti.

#### Materials Availability

This study did not generate new unique reagents.

#### Data and Code Availability

The published article includes all datasets generated and analyzed during this study. Processed bulk RNA-seq data is available online (https://covidgenes.weill.cornell.edu/).

### EXPERIMENTAL MODEL AND SUBJECT DETAILS

#### Human serum and nasopharyngeal swab sample collection for ddPCR

All plasma and nasal swab samples from those with COVID-19 infection, seasonal coronavirus infection, and controls were collected from inpatients at the University of Maryland Medical Center, in Baltimore, USA, between March and May of 2020. Sample collection obtained through informed consent waiver, which was approved by the University of Maryland, Baltimore IRB.

For serum samples, N=10 samples from COVID-19 intubated patients, COVID-19 outpatients, and COVID-19 negative patients were obtained. An equal distribution of N=5 males and females were used for each group. Also, an equal age distribution of patients from 27 to 85 years old was utilized for each group.

For the nasopharyngeal samples the following patient samples were obtained: N=10 SARS-CoV-2 positive patients, N=6 common cold coronavirus samples, and N=6 Coronavirus NL63 samples. For the common cold coronavirus samples the breakdown was the following for the specific viruses: N=2 Coronavirus 229E, Coronavirus HKU1, and N=2 Coronavirus OC43.

#### Human nasopharyngeal swab sample collection for RNA-seq analysis

Patient specimens were processed as described in Butler et al., 2020 (Butler et al., 2021). Briefly, nasopharyngeal swabs were collected using the BD Universal Viral Transport Media system (Becton, Dickinson and Company, Franklin Lakes, NJ) from symptomatic patients. Total Nucleic Acid (TNA) was extracted from using automated nucleic acid extraction on the QIAsymphony and the DSP Virus/Pathogen Mini Kit (Qiagen).

#### Human autopsy tissue collection for RNA-seq analysis

The full methods of the patient sample collection from the autopsy patients are currently available in the Park et al. (Park et al., 2021). All autopsies are performed with consent of next of kin and permission for retention and research use of tissue. Autopsies were performed in a negative pressure room with protective equipment including N-95 masks; brain and bone were not obtained for safety reasons. All fresh tissues were procured prior to fixation and directly into Trizol for downstream RNA extraction. Tissues were collected from lung, liver, lymph nodes, kidney, and the heart as consent permitted. For GeoMx, RNAscope, trichrome and histology tissue sections were fixed in 10% neutral buffered formalin for 48 hours before processing and sectioning. These cases had a post-mortem interval of less than 48 hours. For bulk RNA-seq tissues, post-mortem intervals ranged from less than 24 hours to 72 hours (with 2 exceptions - one at 4 and one at 7 days - but passing RNA quality metrics) with an average of 2.5 days. All deceased patient remains were refrigerated at 4°C prior to autopsy performance.

#### Human urine sample collection

Urine was collected from patients and volunteers at the University of North Carolina at Chapel Hill. All patients provided informed consent prior to participation in IRB-approved research pro-tocols (UNC IRB: 20-0822 [RHS] and 20-0792 [NMB]). Mid-stream urine of outpatients and non-critically ill patients was collected by the clean catch method. Urine of intubated critically ill patients was collected from a port on the Foley catheter. Urine was aliquoted into 5 ml aliquots and stored at -80°C.

Urine aliquots were thawed, and microRNA was extracted from 1 ml per sample using Norgen Urine microRNA Purification Kit (Cat. 29000). Microalbumin and creatinine levels were assessed using Microalbumin 2-1 Combo strips (CLIAwaived Inc, cat# URS-2M).

#### Human peripheral blood mononuclear cells collection for ddPCR

Peripheral blood mononuclear cells (PBMC) were obtained from buffy coats (Hematology and Hemotherapy Center of the University of Campinas) from healthy donors provided Study was approved by the Brazilian Committee for Ethics in Human Studies (CAEE: 31622420.0.0000.5404). PBMCs were isolated as described in Codo et al. (Codo et al., 2020).

#### Cell lines used for miR-2392 mimic experiments

Human SH-SY5Y cells were obtained from the ATCC and grown in Minimum Essential Medium (Gibco) / 10% FBS (Invitrogen) /1% MEM Non Essential Amino Acids (Gibco) / 1%GlutaMAX -l (Gibco). Cells were plated in 3.5 cm dishes and incubated with miR-2392 or control lentivirus particles (MOI 1) for 48h. Cells were harvested and lysed in Trizol reagent and RNA was extracted following manufacturers protocol (Invitrogen).

#### COVID-19 hamster model

Male Syrian hamsters 6-8 weeks old were utilized for efficacy studies with anti-miR-2392 Nanoligomers treatment. Three hamsters were used for each experimental group for a total of 30 hamsters with 10 treatment groups. Hamsters were infected with 10^5^ pfu of the WA01/2020 strain of SARS-CoV-2 passaged twice in Vero E6 cells from the original isolate obtained from BEI Resources. There were 5 major treatment groups (N=6 per group) with two endpoints at day 3 or 7 post-viral challenge (N=3 per endpoint). Groups 1 and 3 were given the Nanoligomer treatment by Intraperitoneal (IP) injection while groups 2 and 4 were given by Intranasal (IN) instillation under ketamine-xylazine anesthesia. Groups 1 and 2 were given single Nanoligomer treatment 24 hours before viral challenge. Groups 3 and 4 were given two doses of Nanoligomer at 24 hours before and 24 hours after viral challenge. The control group 5 was treated with PBS 24 hours prior to and 24 hours following viral challenge by IN instillation. A primary focus for the animal tests was to assess treatment efficacy to reduce viral infection in the pulmonary system. Thus, the Nanoligomers were administered directly to lung tissues via the IN route and the primary endpoints were specifically evaluated for oropharyngeal shedding of virus and lung tissue burden for the virus at necropsy. Injecting the Nanoligomer intravenously (IV) may have also provided a suitable route to target the pulmonary system, however there is concern that the blood filtering organs or uptake in the endothelium may have blunted delivery directly to the lungs.

Treatment efficacy was assessed in multiple ways: 1) Change in daily body weight, 2) oropharyngeal shedding of virus on days 1-3 from all groups post-challenge assayed by plaque assay on Vero E6 cells (PFU/swab), 3) tissue burden of the virus at necropsy on day 3 from 2 lung tubes and turbinates assayed by plaque assay (PFU/100mg), and 4) histopathologic scoring on lungs and turbinates from all hamsters; the histopathological score for individual tissues, inflammation score from the interstitial lung inflammation, and total histopathological scores/assessment was made.

The dose of anti-miR-2392 that was used was calculated to raise blood levels to 10 μM if it were given intravenously. The dose was estimated from the average IC50s determined from our *in vitro* viral screening experiments. The molecular weight of anti-miR-2392 is 15,804. Assuming that hamsters weigh 120 grams and have 8% of body weight as blood, blood volume was approximately 0.01 liters. The dose per hamster was 1.58 mg in a 100 μl volume from an anti-miR-2392 solution.

#### In vitro viral screening model

A549-ACE2 cells, gifted by Dr. Brad Rosenberg (MSSM), were maintained in DMEM (Quality Biological, Gaithersburg, MD; #112-014-101) + 10% Fetal Bovine Serum (Gibco; #26140079) + 1% Penicillin-Streptomycin (Gemini Bio; #400-109). The day prior to treatment, 5,000 A549-ACE2 cells were plated per well in 96-well plates. MiR-2932 was diluted in duplicate in A549-ACE2 media to a starting concentration of 20μM (Run 1) or 22μM (Runs 2 and 3), and then an 8-point 1:2 dilution series was prepared. Media was removed from cells and 90μL of each dilution was transferred to the cells. The plates were incubated for 2 hours at 37°C before being infected with an M.O.I. of 0.1 SARS-CoV-2 WA-1 (provided by Dr. Natalie Thornburg at the Centers for Disease Control and Prevention). Parallel plates were also run and left uninfected to monitor toxicity. Since Runs 2 and 3 were run simultaneously, a single toxicity plate was run for both. All plates were incubated at 37°C for 72 hours before being analyzed via Cell Titer Glo (Promega, Madison, WI; #G7573). Cell viability was compared to uninfected, untreated cells and infected, untreated cells.

### METHOD DETAILS

#### Human peripheral blood mononuclear cell infection with SARS-CoV-2

Monocytes were isolated from PBMCs as described in Toniolo et al. (Toniolo et al., 2015). Cells were infected with SARS-CoV-2 B and P1 lineages as described. B lineage virus was isolated from the second confirmed case in Brazil (GenBank: MT126808.1). P1 was obtained as described by Souza et al. (Souza et al., 2021).

#### miRNA extraction for Droplet Digital PCR (ddPCR)

MiRNA extractions from serum were carried out using the Qiagen miRNeasy serum/plasma kit (#217184). MiRNA extractions from urine samples were carried out using Norgen urine microRNA Purification Kit (Cat. 29000, Norgen Bioteck Corp. Thorold, ON, Canada). Quantitation of miRNA samples was done using a NanoDrop 2000 Spectrophotometer (ThermoFisher Scientific).

#### cDNA generation and ddPCR

First, cDNA was synthesized from miRNA samples using the Qiagen miRCURY LNA RT Kit (Cat. 339340) using a concentration of 5ng/μl for the miRNA per sample. Next, samples were mixed with a 1:20 dilution of the generated cDNA with the BioRad QX200 ddPCR Evagreen Supermix (Cat. 1864034) and the appropriate miRNA primers from miRCURY LNA miRNA PCR Assays (Qiagen). BioRad QX200 Automated Droplet Generator (Cat. 1864101) was used to create emulsion droplets. With the C1000 Touch™ Thermal Cycler with 96–Deep Well Reaction Module (Bio-Rad) the following PCR reaction was used for all the primers: 1 cycle 95°C for 5 min, 40 cycles of 95°C for 30 sec and 58°C for 1 min (the annealing temperature can change depending on the primer), 1 cycle of 4°C for 5 min, and 1 cycle of 90°C for 5 min. Not all miRNA primers sets for ddPCR will have the same annealing temperature, so optimizing the annealing temperature is required for each primer set. Their respective annealing temperatures are found in Table S1. Finally, the QX200™ Droplet Digital™ PCR System (Bio-Rad) quantified the amount of miRNA for each primer set per sample. QuantaSoft software (Bio-Rad) generated the data for each primer set and sample. The same threshold setting was used for all samples per primer set. The concentration (miRNA copies/μl) value generated by QuantaSoft was converted to miRNA copies/ng of serum. These values were used for all miRNA analysis. For all analysis the miRNA concentrations were log_2_(x+1) transformed to allow for easy comparison between miRNAs and samples.

#### Publicly available Bronchial Alveolar Lavage Fluid (BALF) COVID-19 RNA-sequencing data

Fastq files were downloaded from SRA (NCBI BioProject PRJNA605907 (Shen et al., 2020) and NCBI BioProject PRJNA390194 (Ren et al., 2018)). Fastq data files were trimmed using TrimGalore v (0.6.4) with a quality cutoff of 30. Data were then aligned using STAR (v2.7.3) two pass mode to the Human reference genome (GRCh38 v99 downloaded 04-27-2020). Unaligned data were written to a fastq file, and then realigned to the GRCh38 reference genome using Bowtie 2 (v2.3.4.1), and output sam file converted to a bam file using samtools (v1.7). The resultant Bam files were merged, sorted, and read groups added using picard tools (v2.21.3).

#### Publicly available RNA-seq data: A549, Calu-3, NHBE, and COVID-19 lung biopsy

Raw RNA-seq read counts from the publication by Blanco-Melo *et al*. for the A549, Calu-3, and NHBE cell lines as well as post-mortem lung biopsies from two COVID-19 patients were downloaded from the Gene Expression Omnibus (series accession GSE147507) (Blanco-Melo et al., 2020).

#### RNA-seq of Nasopharyngeal Swab COVID-19 patient samples

RNA isolation and library preparation is fully described in Butler, et al. (Butler et al., 2021). Briefly, library preparation on the all nasopharyngeal swab samples’ total nucleic acid (TNA) were treated with DNAse 1 (Zymo Research, Catalog # E1010). Post-DNAse digested samples were then put into the NEBNext rRNA depletion v2 (Human/Mouse/Rat), Ultra II Directional RNA (10 ng), and Unique Dual Index Primer Pairs were used following the vendor protocols from New England Biolabs. Kits were supplied from a single manufacturer lot. Completed libraries were quantified by Qubit or equivalent and run on a Bioanalyzer or equivalent for size determination. Libraries were pooled and sent to the WCM Genomics Core or HudsonAlpha for final quantification by Qubit fluorometer (ThermoFisher Scientific), TapeStation 2200 (Agilent), and qRT-PCR using the Kapa Biosystems Illumina library quantification kit.

#### RNA-seq of COVID-19 autopsy tissue samples

RNA isolation and library preparation is fully described in Park, et al. (Park et al., 2021). Briefly, autopsy tissues were collected from lung, liver, lymph nodes, kidney, and the heart and were placed directly into Trizol, homogenized and then snap frozen in liquid nitrogen. At least after 24 hours these tissue samples were then processed via standard protocols to isolate RNA. New York Genome Center RNA sequencing libraries were prepared using the KAPA Hyper Library Preparation Kit + RiboErase, HMR (Roche) in accordance with manufacturer’s recommendations. Briefly, 50-200ng of Total RNA were used for ribosomal depletion and fragmentation. Depleted RNA underwent first and second strand cDNA synthesis followed by adenylation, and ligation of unique dual indexed adapters. Libraries were amplified using 12 cycles of PCR and cleaned-up by magnetic bead purification. Final libraries were quantified using fluorescent-based assays including PicoGreen (Life Technologies) or Qubit Fluorometer (Invitrogen) and Fragment Analyzer (Advanced Analytics) and sequenced on a NovaSeq 6000 sequencer (v1 chemistry) with 2×150bp targeting 60M reads per sample.

#### miR-2392 mimic experiments in SH-SY5Y cells and RNA-seq

RNA was dissolved in nuclease free water and concentration determined spectrometrically at 260nm using a Biotek plate reader (Biotek). 500ng RNA was used as input for a whole transcriptome library preparation (ThermoFisher Total RNA). Libraries were quantified using a bioanalyzer chip reader (nanoDNA chips: Aglient Technologies) and diluted to 100 pM final concentration. Barcoded libraries were combined and use to seed a OneTouch bead templating reaction (OneTouch2). Cloned libraries were enriched and loaded on 540 Ion Torrent chips. Data were sequenced using the Ion Torrent RNA-seq workflow. Unaligned Bam files were converted to fatsq and aligned to the Grch 38 reference genome using STAR Two pass approach (Dobin et al., 2013) to create gene count tables as described in Overbey et al. (Overbey et al., 2021)(script in supplementary).

#### Anti-miR-2392 Nanoligomer inhibitor design and construction

The Nanoligomer platform was used to design Nanoligomer inhibitors, which are composed of a nanobiohydrd molecule based on antisense peptide nucleic acid (PNA) moiety conjugated to nanoparticle for improved delivery and membrane transport. The PNA moiety was chosen to be 15 bases long (in order to maximize both solubility and specificity), which yielded six potential target sequences within the 20-nucleotide mature human miR-2392. These potential targets were screened using FAST for solubility, self-complementing sequences, and off-targeting within the human genome (GCF_000001405.26) and SARS-CoV-2 viral genome (NC_045512). The antisense sequence complementing miR-2392 nucleotides 2 to 16 (TCTCACCCCCATCCT) was chosen in order to minimize off-targeting while maximizing coverage of the miR-2392 seed region. The Nanoligomer was synthesized (with an N-terminal histidine tag and a 2-(2-(2-aminoethoxy)ethoxy)acetic acid linker) on an Apex 396 peptide synthesizer (AAPPTec, LLC) with solid-phase Fmoc chemistry. Fmoc-PNA monomers were obtained from PolyOrg Inc., with A, C, and G monomers protected with Bhoc groups. Following synthesis, the peptides were conjugated with nanoparticles and purified via size-exclusion filtration. Conjugation and concentration of the purified solution was monitored through measurement of absorbance at 260 nm (for detection of PNA) and 400 nm (for quantification of nanoparticles).

### QUANTIFICATION AND STATISTICAL ANALYSIS

#### Analysis of BALF RNA-seq data

Bam files were imported into Partek Genome Studio v7.0, and gene expression values quantified vs the Grch38 reference annotation guide (Ensembl v99). Samples with fewer than 2 million aligned reads were excluded from further analysis. Genes with fewer than 10 reads in 25% of samples were excluded, and differential gene expression determined using ANOVA with infection status as contrast. Differentially expressed gene files were used in GSEA and IPA to determine biological significance and pathways being regulated.

#### Analysis of Nasopharyngeal Swab RNA-seq data

The nasopharyngeal swab samples were analyzed comparing COVID-19 viral infection to the negative patients and was as previously described in Butler et al. (Butler et al., 2021) and the DESeq2 (Love et al., 2014) was utilized to generate the differential expression data. Heatmaps were displayed using pheatmap (Kolde, 2015). Volcano plots were made use R program EnhancedVolcano (Blighe et al., 2018).

#### Analysis of Autopsy RNA-seq data

The full methods of the analysis from the autopsy patients is currently available in the Park et al. (Park et al., 2021). Briefly, RNA-seq data was processed through the nf-core/rnaseq pipeline (Ewels et al., 2020). This workflow involved adapter trimming using Trim Galore! (https://github.com/FelixKrueger/TrimGalore), read alignment with STAR (Dobin et al., 2013), gene quantification with Salmon (Patro et al., 2017), duplicate read marking with Picard MarkDuplicates (https://github.com/broadinstitute/picard), and transcript quantification with StringTie (Kovaka et al., 2019). Other quality control measures included RSeQC, Qualimap, and dupRadar. Alignment was performed using the GRCh38 build native to nf-core and annotation was performed using Gencode Human Release 33 (GRCH38.p13). FeatureCounts reads were normalized using variance-stabilizing transform (vst) in DESeq2 package in R for visualization purposes in log-scale (Love et al., 2014). Differential expression of genes were calculated by DESeq2. Differential expression comparisons were done as either COVID+ cases versus COVID-controls for each tissue specifically, correcting for sequencing batches with a covariate where applicable, or pairwise comparison of viral levels from the lung as determined by nCounter data. Volcano plots were made use R program EnhancedVolcano (Blighe et al., 2018).

#### Analysis Combining Autopsy and Nasopharyngeal Swab RNA-seq data

To combine the results from the autopsy and nasopharyngeal swab RNA-seq data, we utilized the t-score values from the DESeq2 analysis. Heatmaps were displayed using pheatmap (Kolde, 2015).

#### Gene Set Enrichment Analysis (GSEA)

For pathway analysis on the miR-2392 targets (**Fig. 3**) we utilized ShinyGO (Ge et al., 2020) to determine the significantly regulated pathways for each main cluster in the heatmap. The clustering was determined through k-mean statistics.

For pathway analysis on the miR-2392 mimic RNA-seq data, we utilized fast Gene Set Enrichment Analysis (fGSEA) (Korotkevich et al., 2021). Pathway analysis was done comparing miR-2392 mimics to all controls and the ranked list of genes were defined by the t-score statistics. The statistical significance was determined by 1000 permutations of the genesets (Subramanian et al., 2005).

#### Analysis of proteomic and transcriptomic blood datasets from COVID-19 patients

For the analysis of the miR-2392 targets in the blood tissue, we downloaded whole blood transcriptome data and plasma proteome data from The COVIDome Explorer Researcher Portal (Sullivan et al., 2021). For Transcriptome data we used the following filters: Category “Effect of COVID-19 status”, Platform “Blood”, Statistical test “Student’s t-test”, Adjustment method “none”, Sex “male” and “female”, Age Group “All”. For Proteome data we used the following filters: Category “Effect of COVID-19 status”, Platform “SOMAscan”, Statistical test “Student’s t-test”, Adjustment method “none”, Sex “male” and “female”, Age Group “All”. We created the list of the intersecting genes from both datasets. We analyzed the list using RStudio Desktop 1.3.1093 (RStudio Team (2020). RStudio: Integrated Development Environment for R. RStudio, PBC, Boston, MA URL http://www.rstudio.com/), and visualized data using ggplot2 version 3.3.2 (Wickham, 2016) and ggrepel version 0.8.2 (https://ggrepel.slowkow.com/).

#### Analysis of Monocyte RNA-seq data

The monocyte COVID-19 RNA-Seq data, published under the accession GSE159678 (Rother et al., 2020), was downloaded from SRA and gene expression was quantified using Salmon’s selective alignment approach (Srivastava et al., 2020). The RNA-Seq processing pipeline was implemented using pyrpipe (Singh et al., 2021) (https://github.com/urmi-21/pyrpipe/tree/master/case_studies/Covid_RNA-Seq). Exploratory data analysis and differential expression analysis were performed using MetaOmGraph (Singh et al., 2020).

#### Analysis of A549, Calu-3, NHBE, and COVID lung biopsy data

Each data series was normalized and filtered for low-expressed genes (counts<1). Cell culture samples treated with SARS-CoV-2 were compared to untreated controls and COVID-19-positive patient samples were compared to healthy lung biopsies. Differentially expressed genes were determined from the R-program Limma-Voom (Ritchie et al., 2015) using a linear model with weighted least squares for each gene and the false discovery rate adjusted p-values were calculated.

#### Analysis of miR-2392 mimic RNA-seq data

Differential gene expression was determined using LIMMA-voom (Ritchie et al., 2015). Data were filtered to ensure data contained at least 5 million aligned reads, and average gene counts of > 10. Cell treatments we used as contrasts for differentially expressed gene calculations. These results were then uploaded to GSEA for further analysis. (R script in supplementary section)

#### Conservation of miR-2392 between species

To determined conservation of miR-2392 among different species we utilized UCSC Genome Browser (Kent et al., 2002). Hsa-miR-2392 was entered as input and target species were chosen to include common models for SARS-CoV-2 *in vivo* studies (e.g. mice, ferrets, and hamsters) as well as primates and other animals to provide a wide spectrum of species to observe conservation of miR-2392. The USCS Genome Browser provides the host gene for miR-2392 (i.e. MEG3) and redirects to GTEx (Consortium, 2020) to provide a plot of MEG3 levels based on RNA-seq data on normal tissues.

#### Mapping miR-2392 sequence to SARS-CoV-2 sequences

To explore potential binding sites for miR-2392 we used miRanda software (Enright et al., 2003) to identify all potential binding sites with respect to the SARS-CoV-2 reference genome (Wuhan-Hu-1; NC045512.2) and representative genomes from lineages of concern. The lineages of concern were selected from Global Initiative on Sharing All Influenza Data (GISAID) with each lineage being represented by 14 recent genomes.

#### In silico predictions of genes from miRNAs

Through the use of a Cytoscape (Shannon et al., 2003) plug-in called ClueGo/CluePedia (Bindea et al., 2009), we were able to predict genes targeted by the miRNAs determined. This involved entering all miRNAs in ClueGo and allowing the software to determine the top 50 genes that were significantly regulated and connected to the miRNAs. The predictions only reflect the functions that will be regulated by the miRNAs and do not show whether the function will be activated or inhibited. Lastly, miRNet 2.0 was utilized to predict the diseases and pathways that are associated with the miRNAs (Chang et al., 2020). This was plotted as a dot plot utilizing the R-program ggplot2 (v3.2.1) (Wickham, 2016).

#### ddPCR analysis of miRNA levels in patient samples

First, we normalized the amount of each miRNA measured per body location (nasal, serum, and urine) using the general logarithm transformation. We compared miRNA levels in samples from patients either positive or negative for SARS-nCoV-2 using the student’s t-test (unadjusted) as well as controlling for sex and age using least squares adjustment. Next, we generated receiver operating characteristic curves from each body location to show the performance of a classification model (SARS-nCoV-2 positive versus negative) at all classification thresholds using the absolute, non-transformed levels (miRNA copies per 5 ng RNA). Finally, we performed a subanalysis on each location to compare the variance of each miRNA in SARS-nCoV-2 negative patients compared to other patient groups. For serum and nasal samples, 1-way ANOVA was used to identify variation associated with the patient classification. For urine samples, 2-way ANOVA was used with location (outpatient versus inpatient) and SARS-nCoV-2 positivity as the main factors. If the ANOVA yielded a P < 0.05, Dunnett’s post-test was used to compare subgroup means to the negative patient sample mean.

#### Computational drug repositioning model

Using the SM2miR database (Lui et al., 2007), we assembled an *n* × *m* binary matrix (*X*) containing 3,593 associations between small molecules (*n = 213*, rows) and miRNAs (*m = 1,519*, columns). Each matrix entry (*X_ij_*) was assigned a value of 1 where a small molecule is known to be associated to miRNA, and was 0 otherwise. The chemical notation as a simplified molecular input line entry system (SMILES) was obtained for each small molecule from PubChem. We then calculated the 2D Tanimoto chemical similarity between pairs of small molecules using the MACCS key binary fingerprints with RDKit (RDKit: Open-source cheminformatics; http://www.rdkit.org). Similarly, for each miRNA, we obtained its sequence from miRbase (Kozomara et al., 2019) and computed sequence similarity between miRNAs as the score of their Needleman-Wunsch alignment. We used the binary matrix, together with the chemical and sequence similarities, as input to our state-of-the-art drug target prediction model to predict missing associations in X (Galeano et al., 2021). The model parameters where: p = ½, β_Chem_ = 1, and α_seq_ = 0. To assess the prediction performance of the model, we performed ten-fold cross-validation simulations.

## Notes

### Competing Interest Statement

The authors have declared no competing interest.

### Summary of Updates

This version of the manuscript has been revised to reflect the revisions made after addressing the reviewers comments and resubmitting the paper.

