## Supplemental Figures and Material for "The Great Deceiver: miR-2392’s Hidden Role in Driving SARS-CoV-2 Infection"

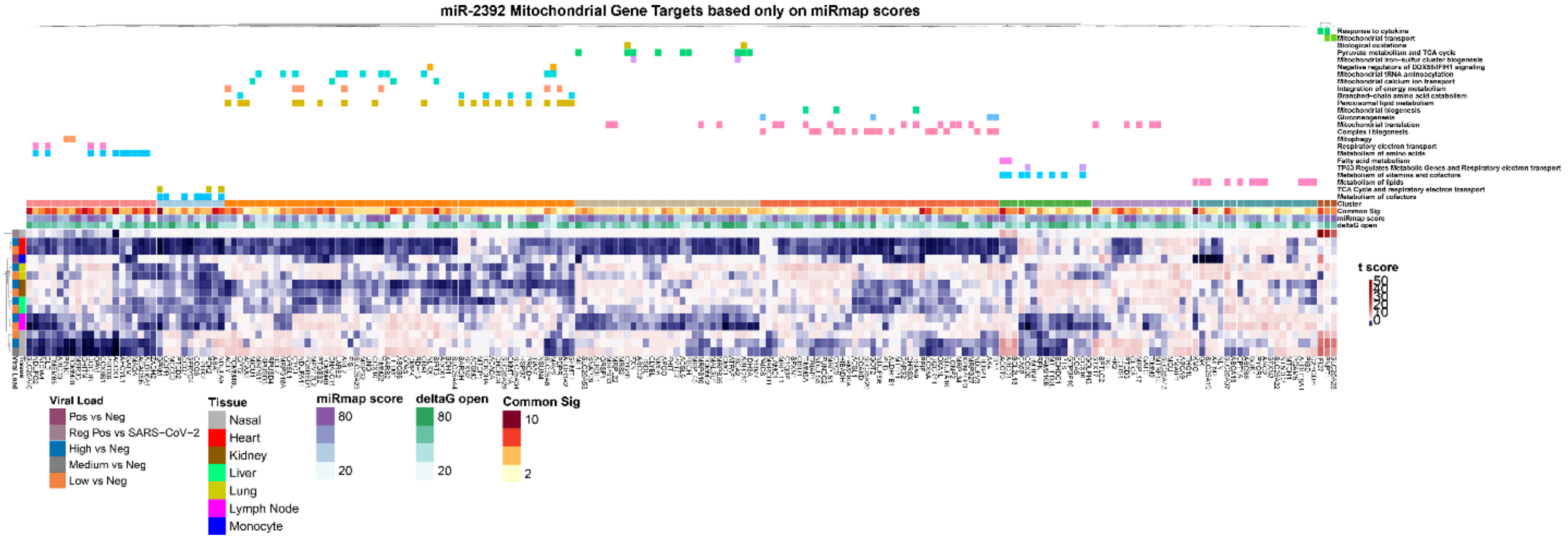


**Figure S1. Mitochondrial gene targets of miR-2392 and regulated pathways. Related to Figure 3.** Differential gene expression analysis for all miR-2392 mitochondrial gene targets significantly expressed in nasopharyngeal swab and autopsy COVID-19 patient tissues. The heatmaps display the t-score statistics for comparing viral load vs negative patient sample for all samples. Main gene clusters were determined through k-mean clustering. Nine main gene clusters were determined and ShinyGO (Ge et al., 2020) was utilized to determine the pathways for each cluster which are displayed on the top panel of the heatmap. Differentially expressed genes are shown with at least one comparison demonstrating a significant adjusted p-value (FDR<0.05) when comparing COVID-19 patients (high, medium or low viral loads) to non-infected control patients (none). Mir-2392 gene targets only determined from miRmap.

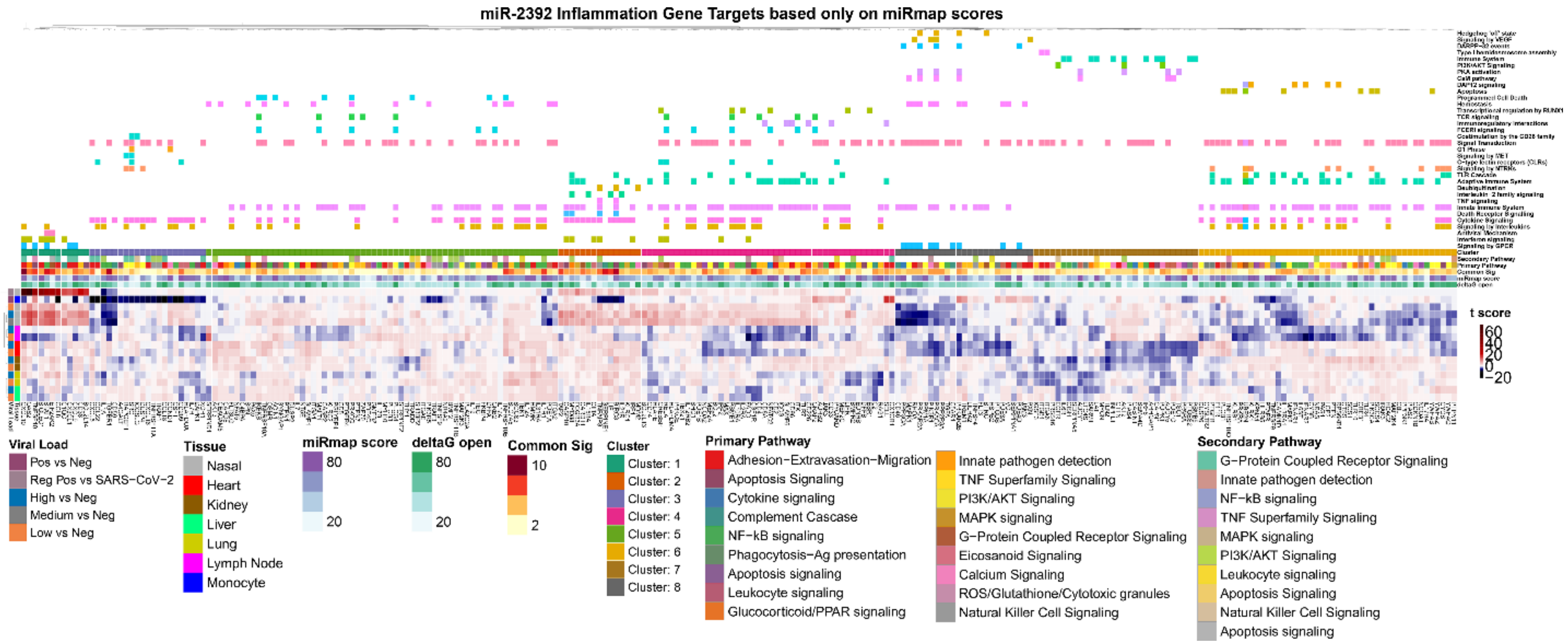


**Figure S2. Inflammatory gene targets of miR-2392 and regulated pathways. Related to Figure 3.** Differential gene expression analysis for all miR-2392 inflammatory gene targets significantly expressed in nasopharyngeal swab and autopsy COVID-19 patient tissues. The heatmaps display the t-score statistics for comparing viral load vs negative patient sample for all samples. Main gene clusters were determined through k-mean clustering. Eight main gene clusters were determined and ShinyGO (Ge et al., 2020) was utilized to determine the pathways for each cluster which are displayed on the top panel of the heatmap. Differentially expressed genes are shown with at least one comparison demonstrating a significant adjusted p-value (FDR<0.05) when comparing COVID-19 patients (high, medium or low viral loads) to non-infected control patients (none). Mir-2392 gene targets only determined from miRmap.

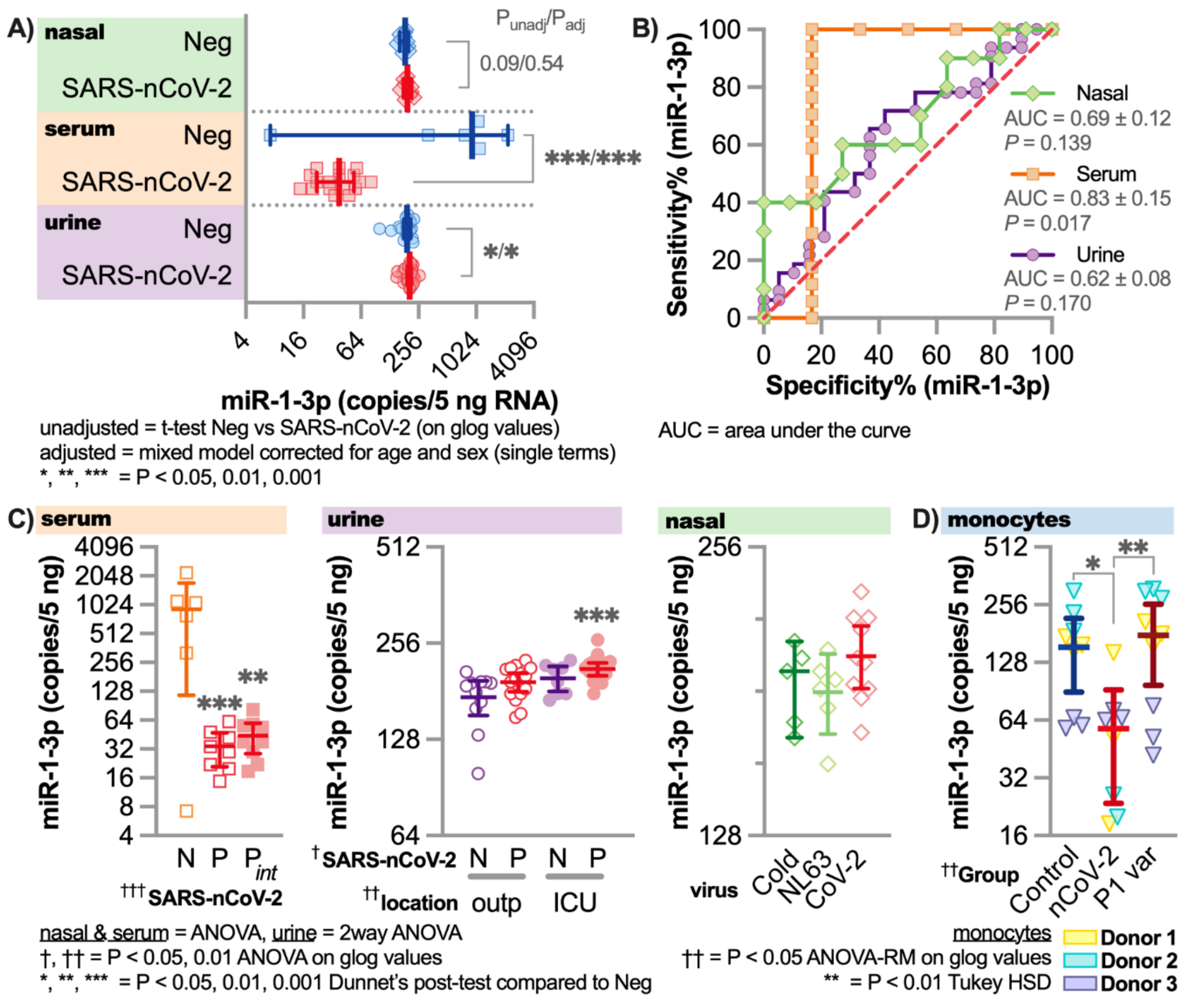


**Figure S3. Circulating miR-1-3p with COVID-19 patients compared to COVID-19 negative patients. Related to Figure 5.** Droplet digital PCR (ddPCR) with specific primer for miR-1-3p was performed on serum, urine, and nasopharyngeal swab samples (including other seasonal coronavirus samples) from COVID-19 positive and negative patients. The miRNA concentration are reported as copies/5ng RNA. **A)** The levels of miRNA-2392 in all tissues from patients grouped as SARS-CoV-2 positive (SARS-nCoV-2) or negative (neg). Unadjusted t-tests comparing the SARS-CoV-2 positive to neg for each tissue are provided and also adjusted statistics comparing the groups with a mixed model corrected for age and sex is provided. **B)** Receiver Operating Characteristic (ROC) curve is provided for miR-1-3p for each tissue comparing SARS-CoV-2 positive to negative patients. **C)** Comparing specific categories within each tissue type between COVID-19 positive and negative patients. N = COVID-19 Negative, P = COVID-19 positive, $P_{int}$ = intubated patients, outp = outpatient, ICU = Intensive care unit/inpatient, Cold = Coronaviruses related to the common cold, NL63 = NL63 coronavirus, and CoV-2 = SARS-CoV-2. **D)** The levels of miR-1-3p in monocytes from healthy donors (n = 3) that were infected with SARS-CoV-2 reference strain and P1 variant. Triplicate conditions were done for each donor. For all plots * = p < 0.05, ** = p < 0.01, and *** = p < 0.001.

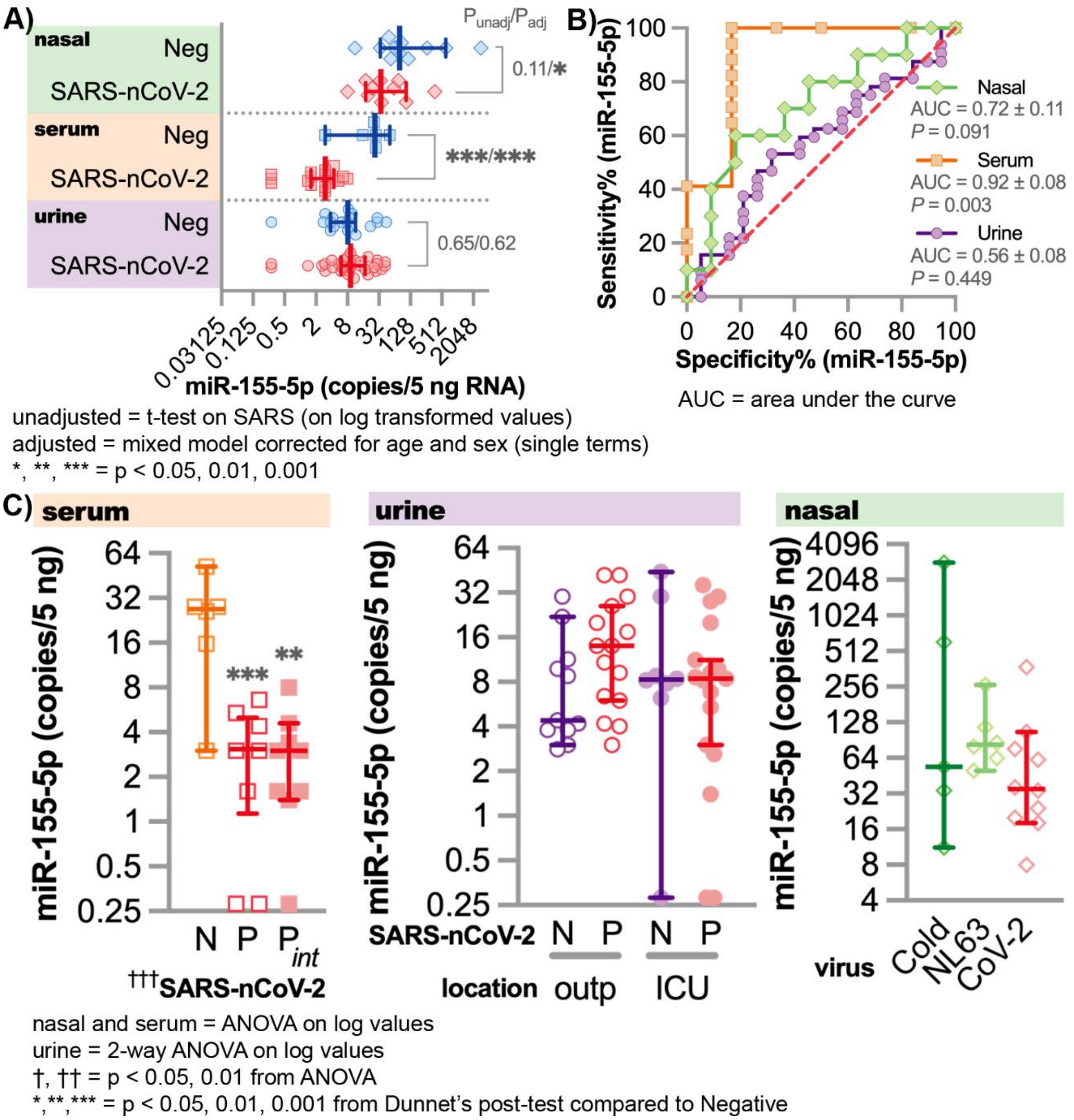


**Figure S4. Circulating miR-155-5p with COVID-19 patients compared to COVID-19 negative patients. Related to Figure 5.** Droplet digital PCR (ddPCR) with specific primer for miR-155-5p was performed on serum, urine, and nasopharyngeal swab samples (including other seasonal coronavirus samples) from COVID-19 positive and negative patients. The miRNA concentration are reported as copies/5ng RNA. **A)** The levels of miRNA-2392 in all tissues from patients grouped as SARS-CoV-2 positive (SARS-nCoV-2) or negative (neg). Unadjusted t-tests comparing the SARS-CoV-2 positive to neg for each tissue are provided and also adjusted statistics comparing the groups with a mixed model corrected for age and sex is provided. **B)** Receiver Operating Characteristic (ROC) curve is provided for miR-155-5p for each tissue comparing SARS-CoV-2 positive to negative patients. **C)** Comparing specific categories within each tissue type between COVID-19 positive and negative patients. N = COVID-19 Negative, P = COVID-19 positive, $P_{int}$ = intubated patients, outp = outpatient, ICU = Intensive care unit/inpatient, Cold

= Coronaviruses related to the common cold, NL63 = NL63 coronavirus, and CoV-2 = SARS-CoV-2. For all plots * = p < 0.05, ** = p < 0.01, and *** = p < 0.001.

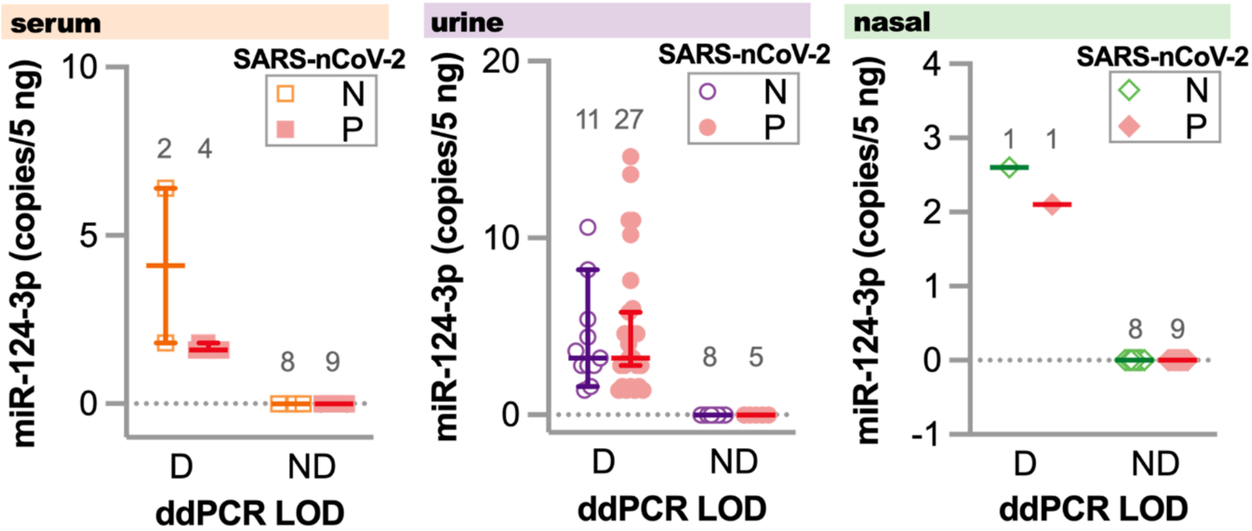


**Figure S5. Circulating miR-124-3p with COVID-19 patients compared to COVID-19 negative patients. Related to Figure 5.** Droplet digital PCR (ddPCR) with specific primer for miR-124-3p was performed on serum, urine, and nasopharyngeal swab samples (including other seasonal coronavirus samples) from COVID-19 positive and negative patients. The miRNA concentration are reported as copies/5ng RNA. For miR-124-3p, the copies/5ng were either equal to 0 or at extremely low levels close to 0 copies/5ng. To try to determine any statistical differences we categorized the groups as ND = Not Determined which are all 0 values or D = Determined which are values > 0 for both N = negative (open symbols) and P = COVID-19 positive patients samples (closed symbols). The number of patients for each column is shown above the points. No significant differences were observed for any of the sample for miR-124-3p.

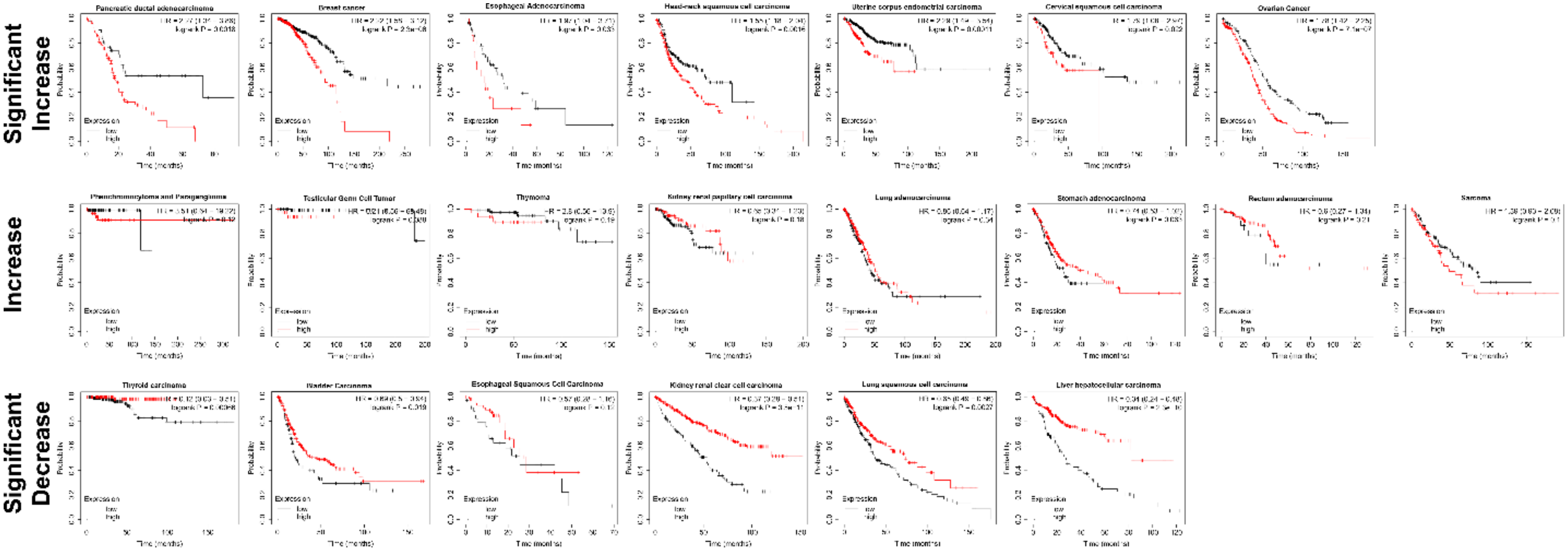


**Figure S6. miR-2392 expression pan-cancer survival analysis. Related to Figure 7.** Kaplan Meier patient survival plots for miR-2392 expression in a pan-cancer analysis was determined utilizing The Kaplan Meier plotter (Nagy et al., 2021). The plots were separated with the top row being cancers which patients had significantly poor survival with high expression of miR-2392, the middle row being cancers which patients had poor survival (but not significant) with high expression of miR-2392, and the bottom row being cancers which patients had significantly better survival with high expression of miR-2392.

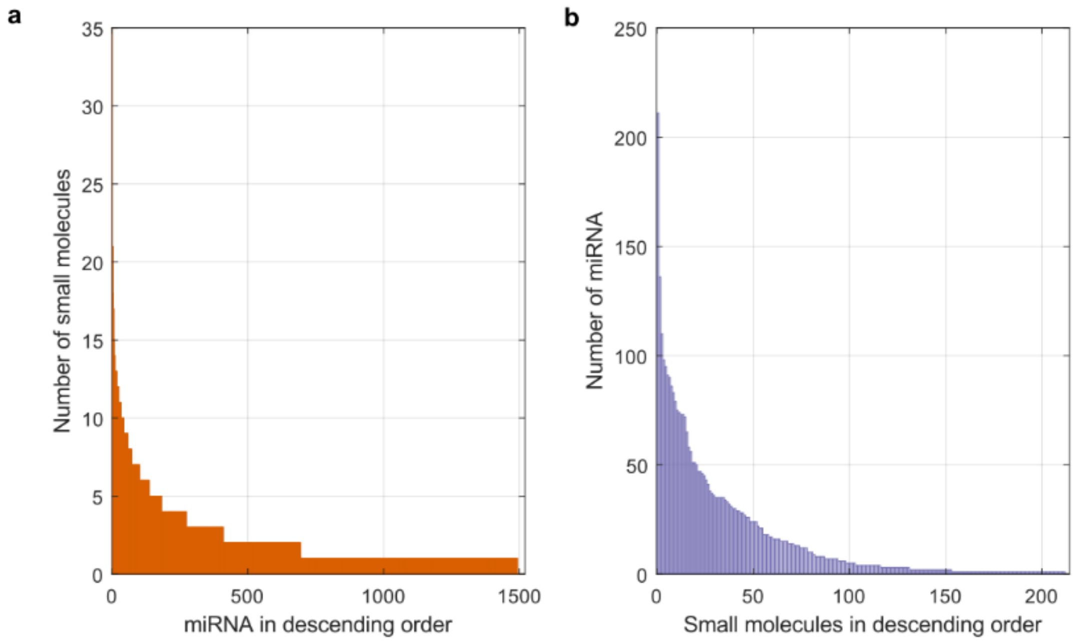


**Figure S7. Small molecules-miRNA dataset statistics Related to Figure 7.** (Left) Number of small molecules associated to miRNAs. (Right) Number of miRNAs associated to small molecules.

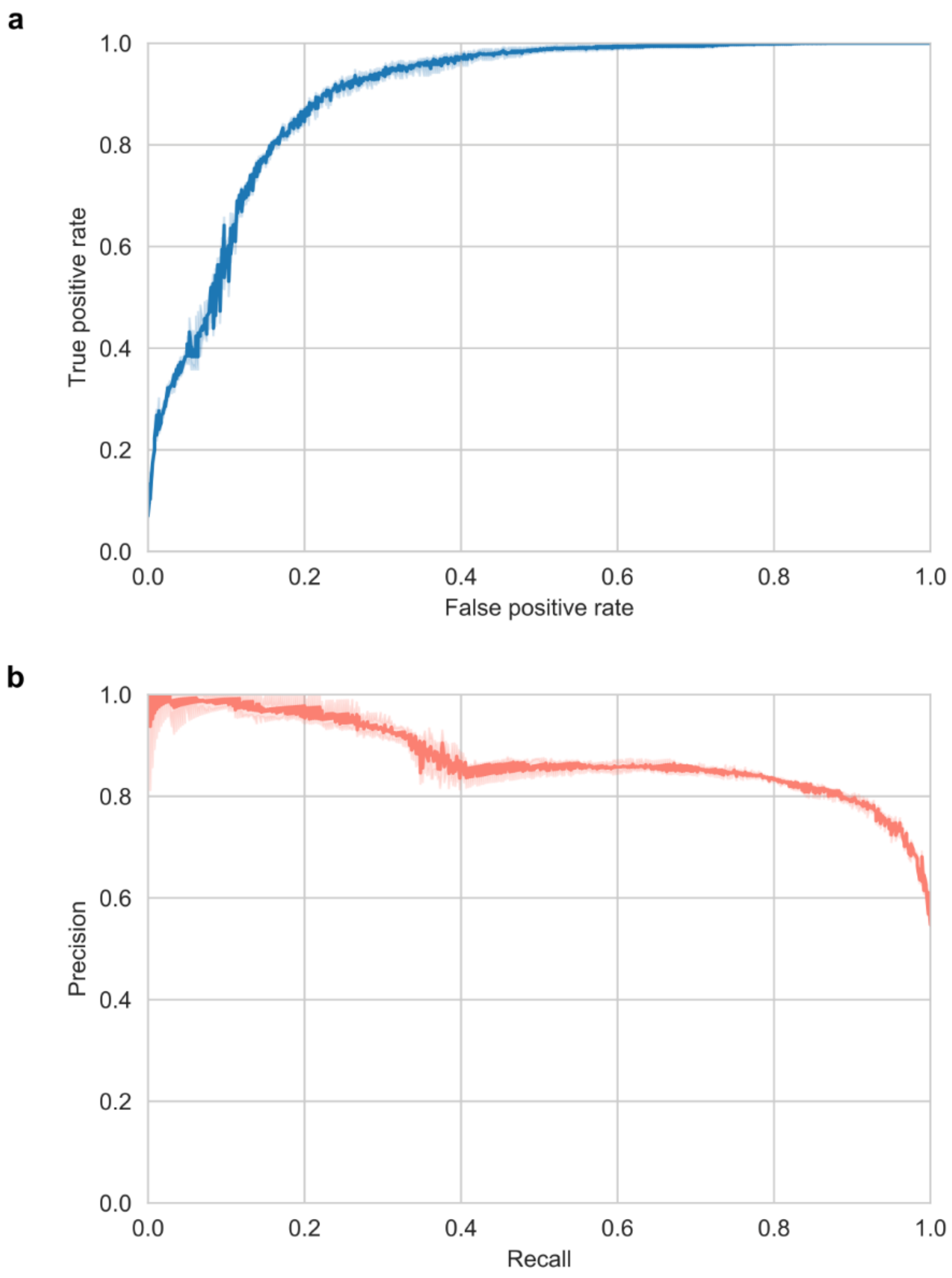


**Figure S8. Performance of our method at predicting missing small molecule-miRNA interactions. Related to Figure 7.** (Top) The mean value of the Receiver Operating Curve (ROC) is shown for a ten-fold cross-validation experiment (dark blue). 95% confidence interval is also shown (light blue). (Bottom) The mean value of the Precision-Recall Curve (PRC) is shown for a ten-fold cross-validation experiment (dark salmon). 95% confidence interval is also shown (light salmon).

a

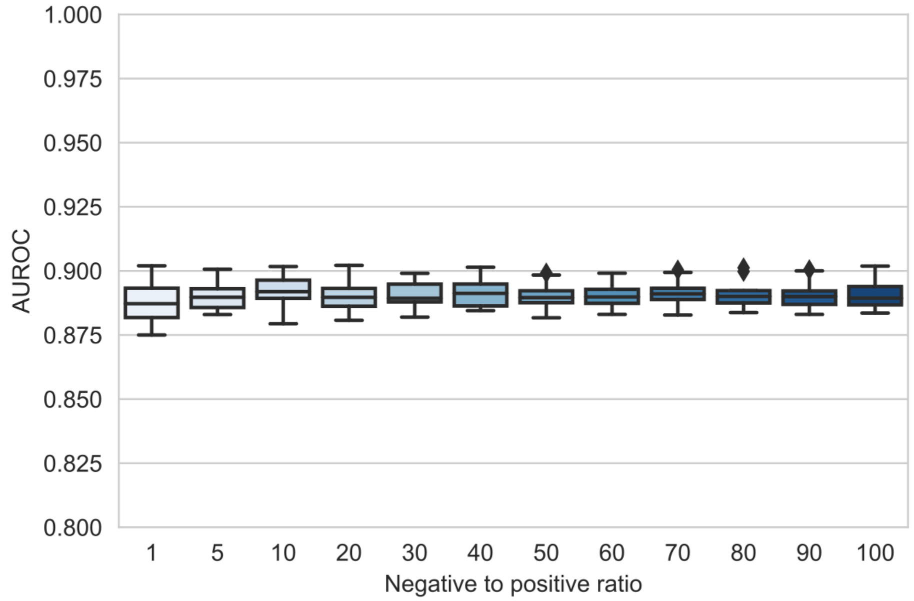


b

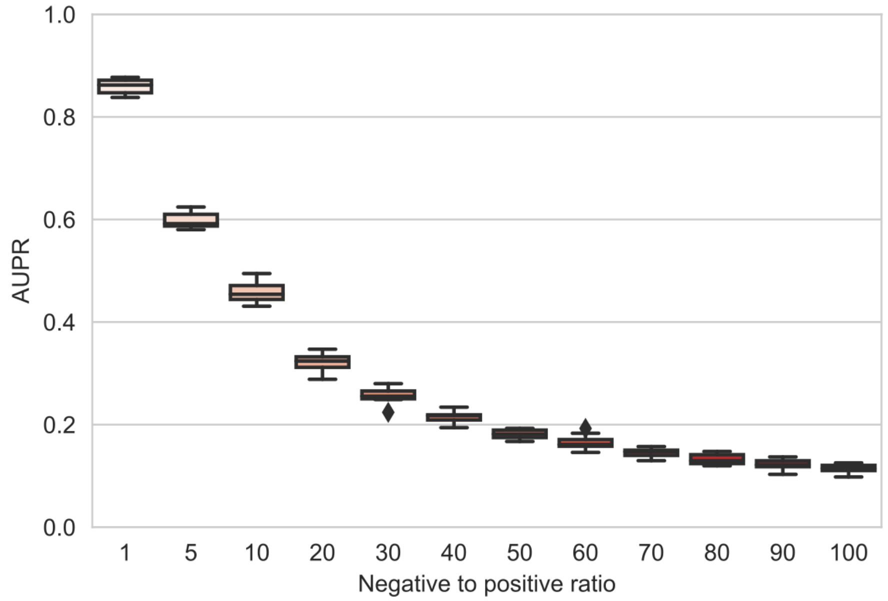


**Figure S9. Performance of our method at predicting missing small molecule-miRNA interactions when controlling for data imbalance. Related to Figure 7.** (Top) Area Under the Receiver Operating Curve (AUROC) was obtained in a ten-fold cross-validation experiment for varying values of the negative to positive label ratio in the test set. (Bottom) Area Under the Precision-Recall Curve (AUROC) was obtained in a ten-fold cross-validation experiment for varying values of the negative to positive label ratio in the test set.

| miRNA target | Primer Annealing temperature | Catalog # |
|---|---|---|
| hsa-miR-2392 | 53°C | Qiagen Cat# YP02104616 |
| hsa-miR-155-5p | 52°C | Qiagen Cat# YP00204308 |
| hsa-miR-124-3p | 58°C | Qiagen Cat# YP00206026 |
| hsa-miR-1-3p | 53°C | Qiagen Cat# YP00204344 |

**Table S1. Annealing temperatures for miRNA primers, related to methods and Figure 5.** Temperatures used for droplet digital PCR to quantify each miRNA target.

**Additional Affiliations:**

**UNC COVID-19 Pathobiology Consortium**

University of North Carolina School of Medicine and Adams School of Dentistry, Chapel Hill, NC 27599, USA

**The UNC COVID-19 Pathobiology Consortium is comprised of:**

Shannon M Wallet[1,2], Robert Maile[2,3], Matthew C Wolfgang[2,4], Robert S Hagan[4,5], Jason R Mock[4,5], Natalie M Bowman[6], Jose L Torres-Castillo[5], Miriya K Love[5], Suzanne L Meinig[4], Will Lovell[1], Colleen Rice[5], Olivia Mitchem[1], Dominique Burgess[1], Jessica Suggs[1], Jordan Jacobs[3]

[1]Department of Oral and Craniofacial Health Sciences, UNC Adams School of Dentistry, University of North Carolina School of Medicine, Chapel Hill, NC, 27599, USA
[2]Department of Microbiology & Immunology, University of North Carolina School of Medicine, Chapel Hill, NC, 27599, USA
[3]Department of Surgery, University of North Carolina School of Medicine, Chapel Hill, NC, 27599, USA
[4]Marsico Lung Institute, University of North Carolina at Chapel Hill, Chapel Hill, NC, 27599, USA
[5]Division of Pulmonary Diseases and Critical Care Medicine, Department of Medicine, University of North Carolina, Chapel Hill, NC 27599, USA
[6]Division of Infectious Disease, School of Medicine, University of North Carolina, Chapel Hill, NC, 27599, USA
